# Preclinical Safety Evaluation of Human Lactoferrin Alpha (Effera^®^)

**DOI:** 10.64898/2026.05.28.728516

**Authors:** Ross Peterson, Nigel Baldwin, Anthony J. Clark, Norbert E. Kaminski, Alan Hoberman, Rajasekhar Pala, Elise Lewis, Andrea Vaden-Harris, Carrie-Anne Malinczak

## Abstract

The preclinical safety of human-equivalent lactoferrin alpha (heqLFα; effera^®^), produced by *Komagataella phaffii*, was evaluated to support its use as a food ingredient in infant formula and products for young children. Genotoxicity was assessed using a bacterial reverse mutation (Ames) assay in five strains of *Salmonella typhimurium* and *Escherichia coli,* and an in vitro micronucleus assay in TK6 cells. Both studies followed OECD guidelines and were conducted up to the recommended limit concentrations (5000 µg/plate and 2000 µg/mL, respectively). A 14-day non-GLP juvenile dose range-finding study and a 13-week GLP juvenile rat toxicity study with a 4-week recovery phase were conducted under ICH S11 guidance. Neonatal Sprague-Dawley rats received heqLFα at 0, 1500, 3000, or 5000 mg/kg body weight/day by twice-daily oral gavage from postnatal day 7 to 98. Bovine lactoferrin (bLF) and whey protein at 5000 mg/kg body weight/day served as reference controls. heqLFα was non-mutagenic and non-clastogenic in both genotoxicity assays. In the 13-week juvenile rat study, no heqLFα-related mortality, clinical signs, developmental, neurobehavioral, or immune toxicity effects were observed. Minor, non-adverse renal findings consistent with high protein intake were observed and largely reversed during recovery. Clinical pathology parameters fully resolved during recovery. Toxicokinetic evaluation showed no systemic accumulation. Based on the absence of toxicologically relevant adverse findings, heqLFα is well tolerated at doses up to 5000 mg/kg body weight/day, establishing a no observed adverse effect level (NOAEL) of 5000 mg/kg/day, the highest dose tested, and supporting its safety across intended populations from birth to adulthood.

## Introduction

Lactoferrin (LF) is an ∼80 kDa iron-binding glycoprotein found in human milk and other biological fluids and mucosal secretions throughout the lifecycle.^1^ It plays a significant role in various physiological processes in early life, including seeding important gut microbes (e.g. bifidobacteria),^2^ immune development,^3^ and iron homeostasis.^4,5^ Given its broad roles and association with improved infant health, particularly in breastfed infants,^6^ LF is considered an appealing food ingredient to be added to infant formula and products for babies and children. To date bovine LF (bLF) has been the only form of LF added to infant formula as a surrogate to human LF (hLF).

Bovine lactoferrin (bLF) is approved as a novel food ingredient for use in infant formula in various markets, including the European Union (EU), China, and the United States. In the United States, bLF is Generally Recognized As Safe (GRAS) for use in milk-based term infant and toddler formulas, as well as in other food applications,^7–10^ indicating scientific consensus on its safety at levels of 100 mg/100g of formula solids. In the EU, bLF has been authorized as a novel food ingredient,^11^ following positive safety opinions from the European Food Safety Authority (EFSA) ^12,13^ to be added to infant and follow-on formula at maximum levels of 100 mg/100 ml (ready-to-drink) and in foods intended for young children at 200 mg/100 g. Clinical studies in infants consuming bLF-fortified formula at concentrations up to 1.0 g/L have consistently demonstrated normal growth, good tolerance, and no reported adverse effects.^4,14–16^

Despite bLF being widely accepted and used in infant formula, the inclusion of hLF would further expand its applicability; however, isolating hLF from human milk (hmLF) for commercial-scale production has ethical and cost challenges that limit feasibility. Human-equivalent lactoferrin (heqLF), such as Helaina heqLF alpha (heqLFα; effera^®^), produced using genetically engineered yeast, *Komagataella phaffii*, can overcome these challenges. Analytical studies confirm that heqLFα is structurally similar to hmLF^17^ and preclinical studies^18,19^ as well as an adult clinical study^20^ demonstrate a strong safety profile for use in non-infant food and dietary supplement application. Other forms of hLF from bioengineered sources have shown safety and tolerance in preclinical animal models^21–28^ and a wide range of vulnerable clinical populations. ^29–33^

From a regulatory perspective most review bodies have a mandated tiered safety assessment approach for food ingredients, requiring at least Tier 1 testing which typically involves genotoxicity studies and a 90-day repeated-dose toxicity study in rodents. Additionally, for new food ingredients targeted at populations which include infants under 16 weeks of age, European Food Safety Agency (EFSA) has issued specific scientific guidance.^34^ A modified 90 day feeding study is required that includes direct oral administration to neonatal/juvenile animals. This age group presents unique physiological, developmental, and nutritional characteristics that require specific considerations for risk assessment. While numerous adult rat toxicology studies have been done on bLF ^35^ and hLF from bioengineered sources^21–24^ and all have shown a no observed adverse effect level (NOAEL) at the highest concentrations tested in studies up to 13 weeks in duration, these studies did not specifically target the juvenile population.

Neonatal/juvenile animal models are well-accepted and have been extensively reviewed by regulatory bodies world-wide to provide key safety data to support new food ingredients in all population groups, specifically including infants less than 16 weeks of age. The most frequently tested ingredients have been the human milk-identical oligosaccharides (HiMO).^36,37^ The typical protocol involves rats fed (e.g., via oral gavage) for 90 days starting from postnatal day (PND) 7 (equivalent to birth in humans) and includes a reference control for comparative safety assessment, and a recovery phase to demonstrate that any adaptive changes are temporary.

In addition to standard general toxicological assessment of LF, because it has well documented immune interactions, immune safety assessments have been recommended by previous expert panels.^38^ In adult rats, heqLFα showed no immunotoxicity risk when T-cell dependent antibody responses and immunophenotyping were evaluated^19^ and low immunogenicity potential in an adult clinical study.

We present here the first in vitro genotoxicity and juvenile rat toxicity studies conducted with heqLFα to assess Tier 1 safety for a novel food ingredient. For the current rat study, the paradigm described above was followed for a 90-day neonatal/juvenile study, with further adaptation to evaluate the toxicokinetics, and potential immunotoxicity to provide further safety reassurance for all intended population groups for heqLFα. The study included high doses of protein (up to 5000 mg/kg body weight (bw)/day); therefore, to account for non-specific protein overload effects, whey protein was included as a comparative control as milk proteins are widely used in infant formula and known to be safe.^39,40^ Additionally, bLF was included as a second high dose reference, due to its functional similarities to hLF and GRAS status in infant formula.^8^ The overall objective was to demonstrate that heqLFα is as safe as whey protein when used as a food ingredient.

## Materials and Methods

### Good Laboratory Practice and Regulatory Compliance

All in vitro genotoxicity assays were executed under Good Laboratory Practices (GLP) regulations^41^ and aligned with Organization for Economic Co-operation and Development (OECD) and Ministry of Health, Labour and Welfare of Japan (MHLW) principles, as well as OECD Test Guidelines (No. 471 for Ames assays, No. 487 for micronucleus assays).^42,43^ Characterization analyses and inspections deviated from GLP (performed per internal SOPs and via process-based inspections), but quality assurance oversight confirmed integrity and compliance throughout. The 13-week neonatal rat toxicity study was also GLP-compliant and followed International Council for Harmonisation (ICH) S11 and S3A guideldines,^41,44,45^ with noted exceptions in test article characterization and immunophenotyping (performed under SOPs but outside GLP). In contrast, the 14-day neonatal rat study was intentionally non-GLP, designed under ICH Tripartite Guideline S11^45^ principles but outside GLP scope as per sponsor strategy.^37,46–48^ As mentioned, this study was designed as an ICH S11-aligned juvenile toxicity study rather than an OECD Test Guideline 408 study.^49^ Although the majority of core repeated-dose toxicology endpoints overlap between OECD 408 and ICH S11-based study designs, the primary distinction is the age and developmental stage of the animals evaluated. OECD 408 is intended for 90-day oral toxicity studies in post-weaning young adult rodents, whereas this study initiated dosing in neonatal/juvenile rats on PND 7 to support assessment during early-life development. This juvenile design is also consistent with EFSA guidance for substances intended for infants below 16 weeks of age, which states that additional studies may be needed for this population and advises that direct dosing of neonatal animals should be considered as soon as possible after birth.^34^

### Test Substances

The primary test substance evaluated across the studies was human equivalent lactoferrin alpha (heqLFα) produced via precision fermentation by *Komagataella phaffii* as described previously.^4,5^ Briefly, heqLFα was purified by microfiltration/diafiltration followed by cation exchange chromatography, then spray-dried into powder form; its purity exceeded 97% on a protein basis and iron saturation was determined to be between 30-60% for all studies reported here. Before use, heqLFα was formulated in 0.9% sodium chloride and adjusted for purity to ensure proper active concentrations and the quantities tested. 0.9% sodium chloride was used as the vehicle control.

In the 14-day and 13-week neonatal rat toxicity studies, bovine lactoferrin (bLF) and whey protein were evaluated as comparative controls since bLF is already approved as an infant formula ingredient and whey protein was used as a high protein control to allow for any dietary imbalance. The bLF (Lactoferrin Co., 21329) contained lactoferrin at a purity of 93.4% to 95.1%. The whey protein control (Hilmar, 200001) was 88.55% protein.

Positive control materials were used in the in vitro genetic toxicology assays. For the bacterial reverse mutation (Ames) assay, specific mutagens were used for different *Salmonella typhimurium* and *Escherichia coli* strains, including 2-nitrofluorene (2NF), sodium azide (SA), ICR-191 (ICR), and 4-nitroquinoline-N-oxide (4NQO) without metabolic activation. 2-aminoanthracene (2AA) served as a positive control in the presence of metabolic activation (**Table 1**). For the in vitro micronucleus assay in TK6 cells, mitomycin C (MMC) was used for 4-hour treatment without metabolic activation, vinblastine sulfate (VIN) for 27-hour treatment without metabolic activation, and cyclophosphamide monohydrate (CP) for 4-hour treatment with metabolic activation. These were chosen as known direct-acting (MMC, VIN) or metabolism-dependent (CP) inducers of micronuclei (**Table 2**). For both in vitro genetic toxicology assays, an exogenous metabolic activation system derived from a phenobarbital/5,6-benzoflavone-induced rat liver microsomal fraction (S9) was used.

**Table 1.** Positive Controls Used in the Bacterial Reverse Mutation (Ames) Assay.

| Strain | Treatment Conditions | Positive Control | CAS No. | Dose |
| --- | --- | --- | --- | --- |
| TA98 | –S9 | 2-Nitrofluorene | 607-57-8 | 2.5 µg/plate |
| TA100 | –S9 | Sodium azide | 26628-22-8 | 1.0 µg/plate |
| TA1535 | –S9 | Sodium azide | 26628-22-8 | 1.0 µg/plate |
| TA1537 | –S9 | ICR-191 | 17070-45-0 | 0.5 µg/plate |
| WP2 uvrA | –S9 | 4-Nitroquinoline-N-oxide | 56-57-5 | 2.0 µg/plate |
| All strains | +S9 | 2-Aminoanthracene | 613-13-8 | 2.5 µg/plate |
| WP2 uvrA | +S9 | 2-Aminoanthracene | 613-13-8 | 10.0 µg/plate |

Abbreviations: –S9, without metabolic activation; +S9, with metabolic activation.

**Table 2.** Positive Controls Used in the In Vitro Micronucleus Assay in TK6 Cells.

| Cell Line | Treatment Conditions | Positive Control | CAS No. | Concentration | Mechanism |
| --- | --- | --- | --- | --- | --- |
| TK6 | 4-hr, –S9 | Mitomycin C | 50-07-7 | 0.0625–0.125 µg/mL | Direct-acting |
| TK6 | 27-hr, –S9 | Vinblastine sulfate | 143-67-9 | 0.0025–0.003 µg/mL | Direct-acting |
| TK6 | 4-hr, +S9 | Cyclophosphamide monohydrate | 6055-19-2 | 11.9 µg/mL | Metabolism-dependent |

Abbreviations: –S9 = without metabolic activation; +S9 = with metabolic activation, TK6 = human lymphoblast cell line.

### Genotoxicity tests

#### Reverse mutation test in Salmonella typhimurium and Escherichia coli (Ames)

In accordance with GLP and the requirements of OECD Test No. 471, the potential for heqLFα to induce reverse gene mutation was examined in *Salmonella typhimurium* strains TA98, TA100, TA1535, and TA1537,^6^ and *Escherichia coli* strain WP2 *uvrA*^7^ in the presence and absence of S9. All were originally purchased from Molecular Toxicology, Inc. (Boone, NC). The Ames assay has been shown to detect various classes of mutagenic chemicals.^7–9^ All heqLFα dose levels (formulated in 0.9% sodium chloride), as well as the positive and vehicle controls, were evaluated in triplicate cultures ±S9 using the preincubation method. heqLFα was initially evaluated in a preliminary dose range-finding assay at 1.00, 5.00, 10.0, 50.0, 100, 500, 1000, and 5000 µg heqLFα/plate in tester strains TA100 and WP2 *uvrA*. heqLFα was freely soluble and noncytotoxic at all dose levels evaluated in strains TA100 ±S9 and WP2 *uvrA* -S9. Cytotoxicity was observed at 5 µg/plate in strain WP2 *uvrA*+S9 however, this response was not dose-responsive and was therefore considered not biologically relevant.

Based upon the results of the dose range-finding assay, heqLFα was evaluated in the definitive mutagenicity assay, in all 5 strains, at 156, 313, 625, 1250, 2500, and 5000 µg/plate (OECD limit dose).

Positive controls for preincubation experiments with and without metabolic activation are in **Table 1**. Vehicle control was 0.9% sodium chloride. Bacterial colonies were counted using Sorcerer Colony Counter (Perspective Instruments, Suffolk, England) and Cyto Study Manager software (Instem Ltd, Suffolk, England).

#### Micronucleus test in cultured human B lymphoblasts (TK6)

In accordance with GLP and the requirements of OECD Test No. 487, the potential of heqLFα to induce micronuclei in human lymphoblasts (TK6 cells) was evaluated, ±S9. This in vitro micronucleus assay detects chromosomal damage (clastogenicity and/or aneugenicity) by measuring cytoplasmic bodies (micronuclei) formed from acentric fragments or whole chromosomes that fail to segregate during anaphase, thereby identifying the clastogenic and aneugenic agents.^10,11^ TK6 cells were originally obtained from Pfizer Global Research and Development (Groton, CT) and subsequently subcloned at the Charles River Testing Facility (Skokie, IL). TK6 stock cultures were maintained in RPMI 1640 (with L-glutamine) supplemented with 10% heat inactivated fetal bovine serum (FBS) and 1% penicillin-streptomycin (Complete Culture Medium, CCM). For experiments, TK6 cells were seeded in CCM at a cell density of 2.50-3.5 × 10^5^ cells/mL and incubated at 36-38 °C and 4-6% CO_2_.

The study involved a range-finding assay and two invalid assay attempts (Trial 1 and Trial 2) due to formulation analysis failures at the high dose followed by Trial 3 as the definitive and accepted study. HeqLFα concentrations tested in the range-finding assays were 3.91 to 2000 µg heqLFα/mL (OECD limit dose). No precipitates or changes in pH were observed at any concentration. Six concentrations were tested (62.5, 125, 250, 500, 1000, and 2000 µg/mL) with the three highest (500, 1000, and 2000 µg/mL) selected for micronucleus evaluation. Cells were exposed to heqLFα at designated doses, as well as to positive controls or 0.9% sodium chloride vehicle alone, -S9 for 4 and 27 hours, and +S9 for 4 h. Positive control compounds are shown in **Table 2**. For the 27-h exposure -S9, cells were harvested at ∼27 h with no further recovery period. For all 4 h treatments (± S9), cultures were allowed a 40-h recovery post-exposure and were harvested at ∼44 h post-initiation. A 40-hour post-treatment recovery period was employed to enhance assay sensitivity compared to the standard 24-hour recovery.^12^ For the definitive assay, duplicate cultures were used, and the three highest concentrations tested were selected for micronucleus evaluation due to the absence of observed cytotoxicity: 500, 1000, and 2000 µg heqLFα/mL.

To confirm that cells underwent sufficient mitotic divisions, as required by OECD Guideline No. 487, cytotoxicity was assessed using cell count data obtained from Coulter counters. Additionally, all cultures were visually examined for signs of cell abnormalities, pH changes (media color), and precipitates.

At harvest, an aliquot of each culture was removed for cell counting. The remaining cells were treated with hypotonic KCl solution (0.28 g KCl per 50 mL deionized water), fixed in methanol:glacial acetic acid, and stored between 2 °C to 8 °C until slide preparation. The final concentrated cell suspension was dropped onto clean glass slides, which were then stained with acridine orange at room temperature. Micronucleus formation was assessed by scoring 1000 cells per duplicate culture (2000 cells per concentration). Slides were randomized and blind coded prior to microscopic analysis. Micronuclei were scored based on: diameter approximately 1/3 or less of the main nucleus, non-refractile, and located in the cytoplasm. Micronuclei were enumerated in slides prepared from three concentrations of the test article, the vehicle control, and a single concentration of the positive control. A Fisher’s Exact 1-tailed test was performed to compare the total number of micronucleated cells in each treatment group against the concurrent vehicle control. Dose-response trend analysis using the Cochran-Armitage test was not warranted, as revertant frequencies across all dose levels of heqLFα approximated vehicle control values in all tester strains, with and without metabolic activation.

### Fourteen-day dose-range finding study in neonatal rats

#### Animals and Housing Conditions

Sixteen Crl:CD® Sprague–Dawley (SD) rat dams with pups randomly distributed into litters at the breeding laboratory (Charles River Laboratories) were delivered to the testing facility 2-4 days postpartum and allowed to acclimate for at least 3 days before experimental procedures commenced. At PND 7, rat pups (weighing 10.4–19.0 g) were allocated into two study phases. The Main Study included 60 pups (30 males, 30 females; six pups/sex/group), and the Extension Phase included 48 pups (24 males, 24 females; six pups/sex/group). Pups were cross-fostered as needed to ensure the appropriate number of male and female pups were assigned to each dose group. Pups remained housed with dams in polycarbonate cages on Bed-o’Cobs® bedding and provided with Crink-l’Nest™, nesting material (The Andersons, Maumee, OH). Environmental conditions were maintained at 21–23 °C and relative humidity of 29–65%, with a 12-hour light/dark cycle. Certified Rodent Diet® #5002 (PMI® Nutrition International LLC, St. Louis, MO) and municipal tap water passed through a reverse osmosis membrane were provided ad libitum. The bedding was changed as often as necessary to keep the animals dry and clean.

#### Test Item Formulation, Administration, and Dosing Schedule

The test article heqLFα, bLF (comparative control), and whey protein (high protein control) were formulated in 0.9% sodium chloride solution. Whey protein was only used in the Extension Phase. A 0.9% sodium chloride solution served as the vehicle control. In the Main Study, heqLFα at 0, 750, 1500, or 3000 mg/kg bw/day, or bLF at 3000 mg/kg bw/day were administered via oral gavage (10 mL/kg). In the Extension Phase, formulations were administered twice daily to achieve a total of 5000 mg/kg bw/day for heqLFα, bLF, or whey protein at a dose volume of 10 mL/kg. Dose formulations were prepared weekly, refrigerated (5 °C), protected from light, and stirred with a stir bar during preparation and administration, and confirmed stable at 0.1–300 mg/mL.

#### Viability and Clinical Signs

Cage-side observations for mortality and moribundity were conducted at least twice daily (morning and afternoon) from the animals’ arrival through the study period. Detailed physical examinations were performed on neonatal rat pups twice weekly during the study period and scheduled on PND 7, 10, 14, and 18, and on the day of scheduled euthanasia. These examinations included skin and fur, eyes, mucous membranes, respiratory and circulatory systems, somatomotor activity and behavior patterns, and feces output and consistency. On each day of dosing observations for clinical signs associated with dosing were conducted before and 1-3 hours following each dose administration.

#### Body Weight and Drinking Water and Food Consumption

Body weight for all neonates selected for dosing was recorded daily from PND 7 to weaning on PND 21. Maternal food and water were monitored and replenished as needed to ensure the health and well-being of the animals.

### 13-week Study of heqLFα Administered by Gavage to Juvenile Rats with a Four-Week Recovery Phase

#### Animals and housing conditions

Sixty-three Crl:CD® (SD) Sprague–Dawley rat dams with random distribution litters (Charles River Laboratories, were delivered to the testing facility 2-4 days postpartum and allowed to acclimate for at least 3 days before experimental procedures commenced. On PND 7, rat pups were allocated into three subsets: Main Study, Recovery Phase, and Toxicokinetic Phase. The Main Study included ten pups/sex/group (**Table 3**); the Recovery Phase (**Table 4)** included five pups/sex/group except Groups 2 and 3 had no Recovery Phase animals; and the Toxicokinetic Phase (**Table 5**) included 9 pups/sex/group in the vehicle control group and 33 pups/sex/group in all other groups. Pups were cross-fostered as needed to ensure the appropriate number of male and female pups were assigned to each group. Pups remained housed with dams in polycarbonate cages on Bed-o’Cobs® bedding and provided with Crink-l’Nest™, nesting material (The Andersons, Maumee, OH).

**Table 3.** Experimental Design (Main Study for Toxicity)

| **Table 3 Experimental Design (Main Study for Toxicity)** |  |  |  |  |  |  |  |
| --- | --- | --- | --- | --- | --- | --- | --- |
| Group | Test Product | Dosing Regimen | Dose Level (mg/kg/d) | Dose Volume (mL/kg) | Dose Concentration (mg/mL) | Males (n) | Females (n) |
| 1 | Vehicle Control | BID | 0 | 10 | 0 | 10 | 10 |
| 2 | heqLFa | BID | 1500 | 10 | 75 | 10 | 10 |
| 3 | heqLFa | BID | 3000 | 10 | 150 | 10 | 10 |
| 4 | heqLFa | BID | 5000 | 10 | 250 | 10 | 10 |
| 5 | Bovine Lactoferrin | BID | 5000 | 10 | 250 | 10 | 10 |
| 6 | Whey Protein Control | BID | 5000 | 10 | 250 | 10 | 10 |

Abbreviations: heqLFa = Human Equivalent Lactoferrin Alpha, BID = Twice Daily

**Table 4.** Experimental Design (Recovery Phase)

| **Table 4 Experimental Design (Recovery Phase)** |  |  |  |  |  |  |  |
| --- | --- | --- | --- | --- | --- | --- | --- |
| Group | Test Product | Dosing Regimen | Dose Level (mg/kg/d) | Dose Volume (mL/kg) | Dose Concentration (mg/mL) | Males (n) | Females (n) |
| 1 | Vehicle Control | BID | 0 | 10 | 0 | 5 | 5 |
| 2 | heqLFa | BID | 1500 | 10 | 75 | - | - |
| 3 | heqLFa | BID | 3000 | 10 | 150 | - | - |
| 4 | heqLFa | BID | 5000 | 10 | 250 | 5 | 5 |
| 5 | Bovine Lactoferrin | BID | 5000 | 10 | 250 | 5 | 5 |
| 6 | Whey Protein Control | BID | 5000 | 10 | 250 | 5 | 5 |

Abbreviations: heqLFa = Human Equivalent Lactoferrin Alpha, BID = Twice Daily, - = Not Applicable

**Table 5.**
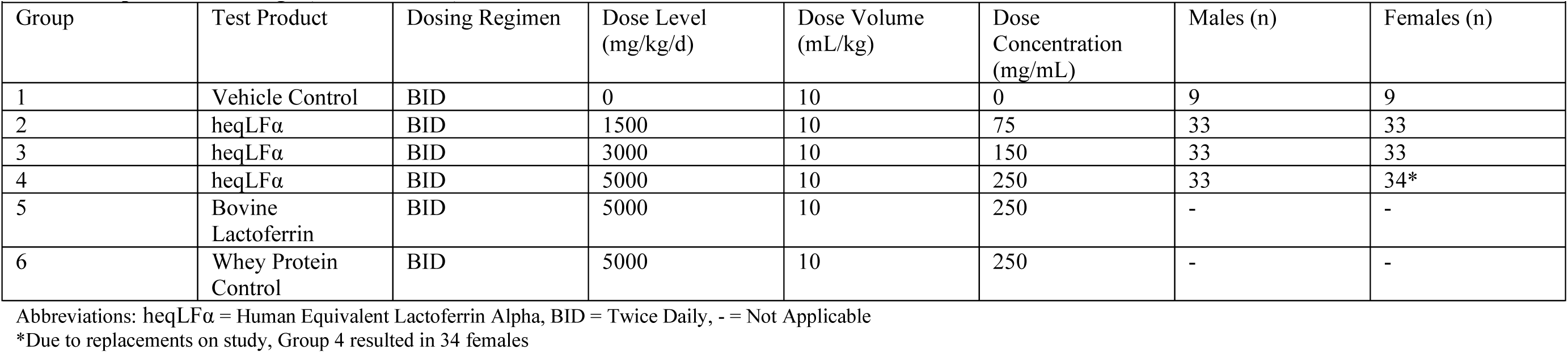
Experimental Design (Toxicokinetics)

| **Table 5 Experimental Design (Toxicokinetics)** |  |  |  |  |  |  |  |
| --- | --- | --- | --- | --- | --- | --- | --- |
| Group | Test Product | Dosing Regimen | Dose Level (mg/kg/d) | Dose Volume (mL/kg) | Dose Concentration (mg/mL) | Males (n) | Females (n) |
| 1 | Vehicle Control | BID | 0 | 10 | 0 | 9 | 9 |
| 2 | heqLFα | BID | 1500 | 10 | 75 | 33 | 33 |
| 3 | heqLFα | BID | 3000 | 10 | 150 | 33 | 33 |
| 4 | heqLFα | BID | 5000 | 10 | 250 | 33 | 34* |
| 5 | Bovine Lactoferrin | BID | 5000 | 10 | 250 | - | - |
| 6 | Whey Protein Control | BID | 5000 | 10 | 250 | - | - |

Abbreviations: heqLFα = Human Equivalent Lactoferrin Alpha, BID = Twice Daily, - = Not Applicable
\*Due to replacements on study, Group 4 resulted in 34 females

Care and use of the animals was in conformity with the American Association for Laboratory Animal Science Policy on the Humane Care and Use of Laboratory Animals. The study protocol and protocol amendments were approved by the Institutional Animal Care and Use Committee (IACUC). Housing was as specified in the USDA Animal Welfare Act (9 CFR, Parts 1, 2, and 3) and as described in the Guide for the Care and Use of Laboratory Animals [8]. The ARRIVE guidelines 2.0 for reporting animal search were followed.

Environmental conditions were maintained at 20–26 °C and relative humidity of 30–70%, with a 12-hour light/12-hour dark photoperiod was maintained, and the rooms were provided with greater than 10 air changes per hour, with 100% fresh air passed through 99.97% HEPA filters. Certified Rodent Diet® #5002 (PMI® Nutrition International LLC, St. Louis, MO) and local water chlorinated after passing through a reverse osmosis membrane were provided ad libitum. The bedding was changed as often as necessary to keep the animals dry and clean. Pups were weaned on PND 21, but no later than PND 28. After weaning, pups were cohoused in same-sex pairs (up to 3 per cage) in solid-bottomed cages by dose group.

#### Test Item Formulation, Administration, and Dosing Schedule

Test animals (Main Study, Recovery Phase, Toxicokinetic Phase) received heqLFα, bLF, whey protein control, or vehicle (0.9 % sodium chloride) at a fixed dose volume of 10 mL/kg, administered via oral gavage twice daily (BID) from PND 7 through PND 98. Formulations were prepared weekly, stored at 5 °C, stirred for ≥30 minutes prior to dosing, and continuously stirred during administration to maintain homogeneity. Group dosing levels were as follows: the vehicle control group received 0 mg/kg bw/day; three heqLFα groups received 1500 (750 BID), 3000 (1500 BID), and 5000 (2500 BID) mg/kg bw/day; and bLF and whey protein control groups each received 5000 (2500 BID) mg/kg bw/day.

#### Animal Observations

##### Clinical signs

Cage-side observations to check for mortality and moribundity were performed twice daily until initiation of dosing and continued on each day of dosing before dose administration and 1-2 hours after dose administration throughout the study period. Cage-side observations involved assessing animals within their cages to monitor for clinical signs, behavioral changes, or reactions to treatment. Detailed clinical observations were conducted outside the cage weekly on PNDs 7, 14, 21, 28, 35, 42, 56, 63, 70, 77, 84, 91, and the day of euthanasia. These assessments included evaluating the general appearance of skin/fur, eyes, mucous membranes, and observing respiratory/circulatory systems, somatomotor activity, and behavior patterns. For the Recovery Phase, these observations were also performed weekly on PNDs 99, 106, 113, 120, 127.

#### Developmental and Reproductive Parameters

Preweaning physical development was assessed in Main Study and Recovery Phase animals. Eye opening was monitored daily beginning on PND 12 until the criterion was achieved or until the day of weaning. The auditory startle reflex and pupil constriction were each evaluated once on PND 21. Post weaning sexual maturation was assessed by daily examination for balano-preputial separation in males beginning on PND 38 and vaginal opening in females beginning on PND 26, continuing until the criterion was achieved for each animal; body weight at the time of sexual maturation was also recorded. Estrous cyclicity was evaluated in females by vaginal lavage cytology beginning 14 consecutive days prior to scheduled euthanasia.

#### Behavioral Assessment

Comprehensive behavioral assessments were conducted during the 13-week testing period in Main Study and Recovery Phase animals to evaluate potential neurobehavioral effects. These assessments included a functional observational battery (FOB) evaluation, automated motor activity, and learning and memory assessment using the Morris water maze. The FOB involved observing animals in their home cage for signs such as posture, convulsions, stereotypy, and tremors, followed by assessments during handling for ease of removal and reactivity, and in an open field to note rearing, gait, vocalizations, respiration, defecation, and general appearance. Further detailed evaluations within the FOB included sensorimotor observations (e.g., touch response, auditory startle, pain response, pupil size) and neuromuscular observations such as body tone and grip strength. Motor activity was assessed by placing rats in individual enclosures to detect and differentiate between fine movements and ambulation over a 60-minute period.

Cognitive function, particularly spatial memory, was examined using the Morris water maze test, conducted over three consecutive days. The first two sessions measured the latency for rats to locate a submerged platform across nine trials each. The third session involved a single probe trial where the platform was removed, and the percentage of time the rat spent in the platform quadrant was recorded to gauge memory retention.

#### Body weight and food consumption

Body weight for all neonates selected for dosing was recorded on each day of dosing from PND 7 and on scheduled euthanasia. Food and water were provided ad libitum, with food consumption quantitatively recorded per cage at least twice weekly from PND 21 for Main Study and Recovery Phases.

#### Terminal Procedures

Organ weights were measured in the Main Study and Recovery Phase animals surviving to scheduled necropsies on PND 99 and PND 127 ± 3 days, respectively, with paired organs weighed together and results expressed both as absolute values and normalized to terminal body and brain weights; and included brain, heart, liver, kidneys, adrenal glands, pituitary, thyroid, thymus, spleen, gonads, and reproductive tract tissues. Organ weights from animals found dead or euthanized in poor condition or in extremis were not recorded, with the exception that testes and epididymides from Recovery Phase males were excised and weighed individually per protocol. Comprehensive histopathology included fixation of cardiovascular, respiratory, gastrointestinal, endocrine, lymphoid, neural, reproductive, and integumentary systems in 10 % neutral buffered formalin, with additional fixatives or special stains employed at the pathologist’s discretion. Microscopic examination of lower-dose groups and Recovery Phase rats was contingent upon test-article–related findings in high-dose Main Study animals. For unscheduled deaths in Main Study and Recovery Phase, gross necropsy and airway perfusion were performed, and tissues retained for possible analysis. No organ weight or histopathology data were collected from Toxicokinetic Phase. Spleens, however, were collected for immunophenotyping analysis.

#### Clinical Pathology

A comprehensive clinical pathology assessment was conducted on blood sampled from the vena cava under deep anesthesia on PND 99 (Main Study) and PND 127 ± 3 days (Recovery Phase animals). Three sample types were collected: hematology (0.3 mL in K₂EDTA), coagulation (0.5 mL in sodium citrate), and clinical chemistry (0.5 mL in serum tubes). Blood for hematology (0.3 mL into K₂EDTA) was maintained at ambient temperature and analyzed within 8 hours of collection or stored at 5 °C and processed within 72 hours. Coagulation samples (0.5 mL into sodium citrate) were centrifuged at room temperature within 30 minutes for ≥15 minutes to yield plasma, which was then snap-frozen on dry ice and held at –70 °C until shipment. Clinical chemistry specimens (0.5 mL in serum separator tubes) were allowed to clot 20–60 minutes at room temperature, centrifuged ≥15 minutes to isolate serum, then frozen on dry ice and stored at –20 °C pending analysis.

Hematology panels included red and white cell counts, hemoglobin, hematocrit, platelet and reticulocyte counts, and differential leukocyte enumeration; coagulation assays measured activated partial thromboplastin time (aPTT), prothrombin time (PT), and fibrinogen; and chemistry analyses included liver enzymes, renal markers, electrolytes, lipids, proteins, glucose, and bilirubin.

#### Toxicokinetic Evaluation

Toxicokinetic (TK) evaluations were conducted on 3 animals/sex/time point on PND 7 and PND 98 to characterize systemic exposure and TK parameters of heqLFα. Blood was drawn via cardiac puncture on PND 7 and via jugular vein (or vena cava during terminal collection) on PND 98 at 0 (predose for only PND 98), 0.5, 1.5, 4, 8.5, 9.5, 12, and 24 hours post-dose. Serum samples collected in serum-separator tubes, held at room temperature during clotting, centrifuged at 5 °C for 10 minutes at 3500 rpm, then aliquoted into duplicate polypropylene tubes, snap-frozen on dry ice, and stored at –70 °C within two hours of collection. Quantitation of heqLFα, was accomplished by validated electrochemiluminescence immunoassay (ECLIA), with incurred sample reanalysis to confirm assay precision and accuracy.

TK parameters were calculated using the linear trapezoidal method with linear interpolation, with concentration values below the limit of quantitation (<1 ng/mL) treated as zero, and included time to maximum concentration (t_max_), maximum observed concentration (C_max_), C_max_/dose, area under the concentration-time curve to the last quantifiable sample (AUCt_last_), AUCt_last_/dose, area under the concentration-time curve from 0 to 24 hours (AUC_0-24hr_), AUC_0-24hr_/dose, time of final quantifiable concentration (t_last_), female-to-male AUC ratio (F:M AUC), female-to-male C_max_ ratio (F:M C_max_), the AUCt_last_ following repeat dosing divided by AUC_tlast_ during the initial dosing interval (RAUC), and the C_max_ following repeat dosing divided by C_max_ during the initial dosing interval (RC_max_).

#### Immunophenotyping

##### Tissue Processing

Animal husbandry, test article administration, and necropsy/organ collection were performed at the Testing Facility, Charles River Laboratories (Horsham, PA). Spleens from the TK animals (N=3 for vehicle control; N=6 for heqLFα (all 3 dose groups), bLF, and whey protein group) were collected, stored in medium, blind-coded prior to shipment, and shipped to the Kaminski laboratory (Michigan State University, East Lansing, MI) at 4 °C within 24 hours of collection. Immunophenotyping analysis was conducted blinded to treatment group and dose level. Splenocytes were isolated by mechanical disruption of the spleen and made into single cell suspensions in RPMI 1640 medium supplemented with 5% fetal calf serum (FCS) and penicillin/streptomycin. Red blood cells were lysed using Zap-oglobin-II Lytic Reagent (Beckman Coulter) prior to counting splenocytes on a Z1 Beckman Coulter Counter.

##### Flow Cytometry

A total of 2 x 10^6^ splenocytes were placed in a 96-well round bottom culture plate for surface and intracellular antibody staining. Splenocytes were washed using Hank’s Balanced Salt Solution (HBSS, (pH 7.4; Invitrogen) and stained with LIVE/DEAD Fixable Near-IR Dead Cell Stain (Gibco Invitrogen) to assess cell viability. Splenocytes were washed with FACS buffer (1× HBSS (pH 7.5) containing 1% BSA and 0.1% sodium azide) and Cell Fc Receptors (FcRs) were blocked with purified mouse anti-rat CD32 (BD Biosciences). After a 15-minute incubation at 4°C splenocytes were stained for surface proteins using the following antibodies. For the identification of Granulocyte/Macrophage/DC and B cells, the following antibodies were used: CD45 (OX-1) (Biolegend), CD172a (clone OX-41) (BD Bioscience), CD11b (clone WT.5) (Biolegend), CD25 (clone OX-39) (Biolegend), and AffiniPure Anti-Rat IgG + IgM (Jackson Immuno Research). For the identification of T cells, the following antibodies were used: CD3 (clone 1F4) (Biolegend), CD4 (clone W3/25) (Biolegend), CD8 (clone OX-8) (Biolegend) and CD25 (clone OX-39) (Biolegend). Splenocytes were incubated at 4°C and washed three times with FACS buffer and fixed with FOXP3/Transcription Factor Staining Buffer Set (Invitrogen) fixative and stored at 4°C until intracellular staining.

To measure intracellular FoxP3, splenocytes were washed and incubated with 1x Permeabilization Buffer Solution (Invitrogen) for 15 minutes, then incubated with anti-rat FoxP3 (Invitrogen) for 30 minutes. Splenocytes were washed 3X with the permeabilization buffer and then washed one more time with FACS buffer and resuspended in FACS buffer. Flow cytometric analysis was performed on a Cytek Northern Lights full spectral analyzer (Cytek Biosciences) and analyzed using FlowJo v10.9.0 (Tree Star, Ashland, OR) software.

#### Statistical Analyses

All statistical analyses were performed using SAS 9.4 (SAS Institute Inc., Cary, NC) with two-sided hypothesis tests at α = 0.05. Clinical and necropsy observations were summarized by sex and group for predefined intervals (PND 7–10, 10–14, 14–17, 17–21, and 7–21 days), reporting descriptive statistics (means, standard deviations or coefficients of variation, ratios, percentages, and incidences). Categorical outcomes were compared by Fisher’s exact test. Continuous endpoints underwent Levene’s test for homogeneity of variances; if variances were equal, an overall one-way ANOVA F-test was applied, otherwise a Kruskal–Wallis test was used. Significant omnibus tests (p ≤ 0.05) were followed by Dunnett’s or Dunn’s multiple comparisons, respectively, against the control group. Morris Water Maze latency data were partitioned into three blocks of three trials per session and analyzed by repeated-measures ANOVA with Group, Time, and Group×Time interaction as fixed effects; the optimal covariance structure (compound symmetry or first-order autoregressive, homogeneous or heterogeneous) was selected by corrected Akaike’s Information Criterion and likelihood ratio tests. Significant group effects triggered Dunnett’s post hoc comparisons across all time points or within individual time points depending on interaction significance. Probe trial percentages were evaluated by one-way ANOVA per sex, with model selection via likelihood ratio tests and Dunnett’s comparisons for significant group effects. Immunophenotyping statistical analyses were performed using GraphPad Prism version 10.0.02 (GraphPad Software, La Jolla, CA). To determine statistically significant changes between the treatment groups and the control (0 mg/kg bw/day) within male and female rats, a one-way ANOVA Dunnett’s multiple comparisons test was used.

## Results

### Bacterial reverse mutation test

No signs of cytotoxicity or mutagenicity were noted in any of the 4 strains of *Salmonella typhimurium* (TA98, TA100, TA1535, and TA1537) or in the *Escherichia coli* strain WP2 *uvrA* compared with vehicle control counts with or without metabolic activation up to the top concentration of 5000 µg heqLFα/plate (**Table 6**). Positive controls, applying 2NF, 4NQO, SA in the absence, and 2AA in the presence of S9, verified the sensitivity of the assay, as at least two- to three-fold increases in the mean revertant colony numbers compared to vehicle control (0.9% sodium chloride) were produced. Thus, the absence of gene mutation following exposure to heqLFα was confirmed.

**Table 6.** Bacterial reverse mutation test performed with heqLFα.

| Test item<br>(mg/plate) | Number of revertant colonies per plate |  |  |  |  |  |  |  |  |  |
| --- | --- | --- | --- | --- | --- | --- | --- | --- | --- | --- |
|  | TA98 |  | TA100 |  | TA1535 |  | TA1537 |  | WP2 *uvrA* |  |
|  | without S9 | with S9 | without S9 | with S9 | without S9 | with S9 | without S9 | with S9 | without S9 | with S9 |
| **Preincubation test** |  |  |  |  |  |  |  |  |  |  |
| Negative control<sup>a</sup> | 14.3 ± 4.0 | 17.0 ± 3.6 | 93.3 ± 7.5 | 106.3 ± 21.3 | 8.7 ± 5.5 | 7.0 ± 2.6 | 5.0 ± 2.0 | 7.3 ± 0.6 | 37.3 ± 1.2 | 46.3 ± 4.5 |
| 156 | 14.7 ± 3.2 | 13.7 ± 5.1 | 90.0 ± 4.0 | 97.3 ± 3.1 | 7.3 ± 3.1 | 7.7 ± 2.1 | 4.7 ± 2.9 | 6.0 ± 1.0 | 47.0 ± 13.7 | 51.0 ± 12.1 |
| 313 | 17.0 ± 6.6 | 20.0 ± 2.0 | 95.7 ± 14.7 | 89.3 ± 13.8 | 9.7 ± 1.5 | 7.3 ± 4.9 | 4.0 ± 1.0 | 4.7 ± 2.5 | 39.3 ± 16.2 | 51.0 ± 5.6 |
| 625 | 20.3 ± 2.1 | 18.0 ± 1.7 | 83.3 ± 6.7 | 81.3 ± 7.4 | 10.7 ± 3.1 | 6.3 ± 1.5 | 5.0 ± 1.7 | 8.3 ± 3.5 | 35.3 ± 1.2 | 54.7 ± 8.5 |
| 1250 | 13.0 ± 2.0 | 22.7 ± 8.0 | 90.3 ± 6.4 | 89.7 ± 12.5 | 7.7 ± 4.9 | 7.7 ± 3.8 | 2.7 ± 1.5 | 4.7 ± 3.1 | 37.0 ± 7.2 | 47.0 ± 10.4 |
| 2500 | 23.7 ± 12.5 | 13.3 ± 6.5 | 86.7 ± 9.3 | 89.3 ± 10.1 | 10.0 ± 5.0 | 9.7 ± 1.5 | 3.7 ± 2.3 | 4.0 ± 1.7 | 39.0 ± 13.5 | 47.0 ± 9.5 |
| 5000 | 17.7 ± 10.0 | 18.0 ± 5.6 | 88.0 ± 6.2 | 94.3 ± 23.7 | 10.3 ± 2.3 | 5.7 ± 1.5 | 3.3 ± 1.5 | 4.3 ± 1.2 | 41.0 ± 10.4 | 37.3 ± 7.2 |
| Positive control<sup>b,c</sup> | 1078.7 ± 71.8 | 3119.3 ± 234.9 | 528.3 ± 89.6 | 3482.7 ± 62.6 | 559.7 ± 31.8 | 324.0 ± 48.1 | 62.0 ± 13.2 | 229.3 ± 22.2 | 1381.7 ± 45.5 | 380.7 ± 48.5 |

Abbreviations: heqLFα = Human Equivalent Lactoferrin, Alpha TA = Typhimurium Strain A, S9 = metabolic activation
<sup>a</sup>Negative control: 0.9% sodium chloride
<sup>b</sup>Positive controls without S9: 2NF (2.5 ug/plate) for TA98, 4NQO (2 ug/plate) for WP2uvrA, ICR (0.5 ug/plate) for TA1537, SA (1 ug/plate) for TA100 and TA1535
<sup>c</sup>Positive controls with S9: 2AA (2.5 μg/plate) for all Salmonella strains; 2AA (10.0 μg/plate) for WP2 uvrA
Values are means (n=3) ± standard deviations.

### In vitro mammalian cell micronucleus test

heqLFα was tested up to the maximum concentration of 2000 µg/mL in TK6 cells both in the absence (4- and 27-hour exposures) and presence (4-hour exposure) of S9 metabolic activation, with negative (0.9 % NaCl) and positive controls (mitomycin C, vinblastine sulfate, cyclophosphamide) included. No cytotoxicity, precipitates, or increases in micronucleated cell frequency were observed, and all frequencies for heqLFα and vehicle controls fell within the laboratory’s historical control range (**Table 7**). In contrast, the positive controls induced significant (p ≤ 0.05) chromosomal damage, confirming assay sensitivity and validity. Accordingly, heqLFα was deemed non-clastogenic and non-aneugenic under these conditions.

**Table 7.** In vitro Micronucleus Test in TK6 Cells Exposed to Human Equivalent Lactoferrin Alpha (heqLFα)

| Test item concentration [μg/mL] | Cytotoxicity (%)<sup>a</sup> | Number of mononucleated cells scored<sup>b</sup> | Total number of micronucleated mononucleated cells<sup>c</sup> |
| --- | --- | --- | --- |
| **4-hour treatment without metabolic activation** |  |  |  |
| Vehicle control (0.9% Sodium Chloride) | 0 | 2000 | 6 |
| Mitomycin C (0.125) | 37.54 | 2000 | 104** |
| heqLFα (500) | 0 | 2000 | 4 |
| heqLFα (1000) | 1.90 | 2000 | 6 |
| heqLFα (2000) | 1.70 | 2000 | 5 |
| **27-hour treatment without metabolic activation** |  |  |  |
| Vehicle control (0.9% Sodium Chloride) | 0 | 2000 | 6 |
| Vinblastine sulfate (0.003) | 43.05 | 2000 | 96** |
| heqLFα (500) | 4.28 | 2000 | 7 |
| heqLFα (1000) | 2.25 | 2000 | 10 |
| heqLFα (2000) | 6.15 | 2000 | 9 |
| **4-hour treatment with metabolic activation** |  |  |  |
| Vehicle control (0.9% Sodium Chloride) | 0 | 2000 | 12 |
| Cyclophosphamide (11.9) | 57.61 | 2000 | 100** |
| heqLFα (500) | 3.48 | 2000 | 7 |
| heqLFα (1000) | 0.78 | 2000 | 5 |
| heqLFα (2000) | 3.04 | 2000 | 3 |

<sup>a</sup> Cytotoxicity values represent the mean percent cytotoxicity based on Relative Population Doubling (RPD), calculated as 100 - RPD.
<sup>b</sup> The OECD Guideline No. 487 recommends scoring 2000 cells per concentration when cytoB is not used. In this study, 2000 mononucleated cells were evaluated per concentration, which corresponds to 1000 cells per culture for the duplicate cultures.
<sup>c</sup> Represents the combined count of micronucleated mononucleated cells from duplicate cultures for each concentration.
\* $p \leq 0.05$ , \*\* $p \leq 0.01$ using Fisher's Exact 1-tailed test, indicating statistically significant difference from the concurrent vehicle control. Statistical significance for positive controls demonstrates the assay's validity.

### Fourteen-day dose range-finding and oral tolerability study in neonatal rats

Three deaths occurred in the bLF 5000 mg/kg bw/day group, 1 in the heqLFα groups at 5000 mg/kg bw/day) and two at 1500 mg/kg bw/day). Four of these deaths had no clear cause of death, one was gavage-related perforation and one pup was cannibalized. The deaths occurred across different groups on non-consecutive days spanning PND 9 to PND 20, with no identifiable pattern related to the test article. No test-article–related clinical signs were observed at doses ≤3000 mg/kg bw/day; at 5000 mg/kg bw/day, sporadic tremors and transient abnormal respiratory sounds were noted but were not considered adverse. Mean body weights and gains were unaffected at ≤3000 mg/kg bw/day, whereas at 5000 mg/kg bw/day both LF groups showed weight gains greater than vehicle control but comparable to the whey protein comparative control. No macroscopic lesions were detected at scheduled necropsy in any group. These findings indicate that 5000 mg/kg bw/day (BID) is tolerated when administered to neonatal rats starting at PND 7 and supports the selection of up to 5000 mg/kg bw/day for definitive neonatal toxicity evaluation to characterize dose-response relationships.

### Repeated dose 13-week oral (gavage) toxicity study in juvenile rats Mortality and clinical observations

In the Main Study and Recovery Phase, excluding intubation- or dosing-related accidents, there were no heqLFα-related deaths. One male in the vehicle control group and three females in the bLF group (5000 mg/kg bw/day) died or were euthanized early. All remaining animals administered heqLFα (1500, 3000, or 5000 mg/kg bw/day), bLF (5000 mg/kg bw/day), or whey protein (5000 mg/kg bw/day) survived to scheduled necropsy.

There were no heqLFα-related clinical observations in males or females throughout the Main Study or Recovery Phase. Clinical signs observed sporadically across groups were consistent with common background findings in juvenile Sprague-Dawley rats and showed no dose-response relationship.

### Body weights, body weight Gain, and Food Consumption

At the end of the dosing phase (PND 99), mean body weights of males and females administered heqLFα at 1500, 3000, or 5000 mg/kg bw/day were comparable to vehicle controls (**Figures 1A and 1B**). Although transient, statistically significant differences in body weights and body weight gains were observed at earlier time points during the study, these were not present at the terminal evaluation and did not demonstrate dose dependency and were considered non-adverse.

**Figure 1.**
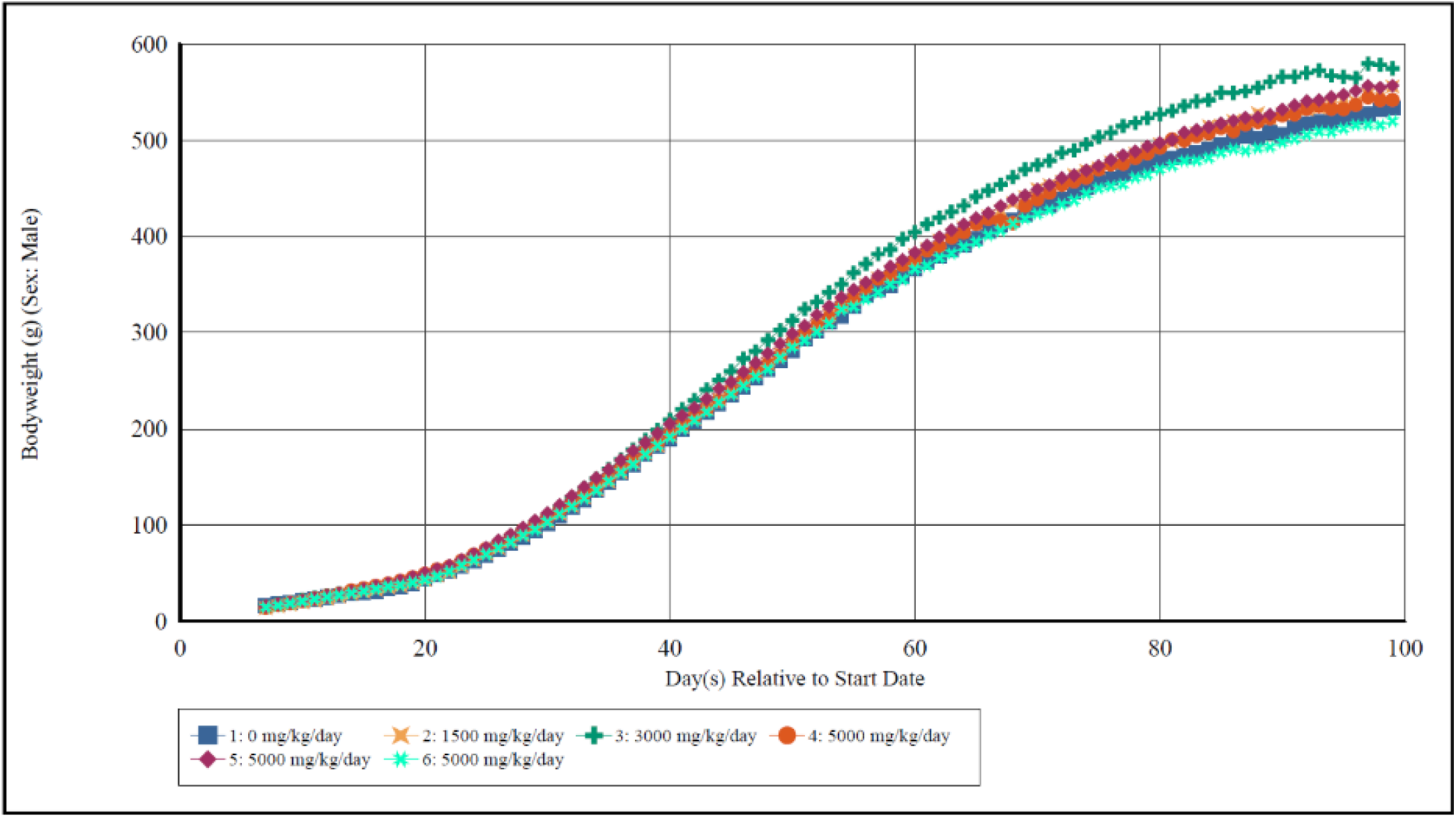
A. Summary of Male Group Mean Body Weight

**Figure 1.**
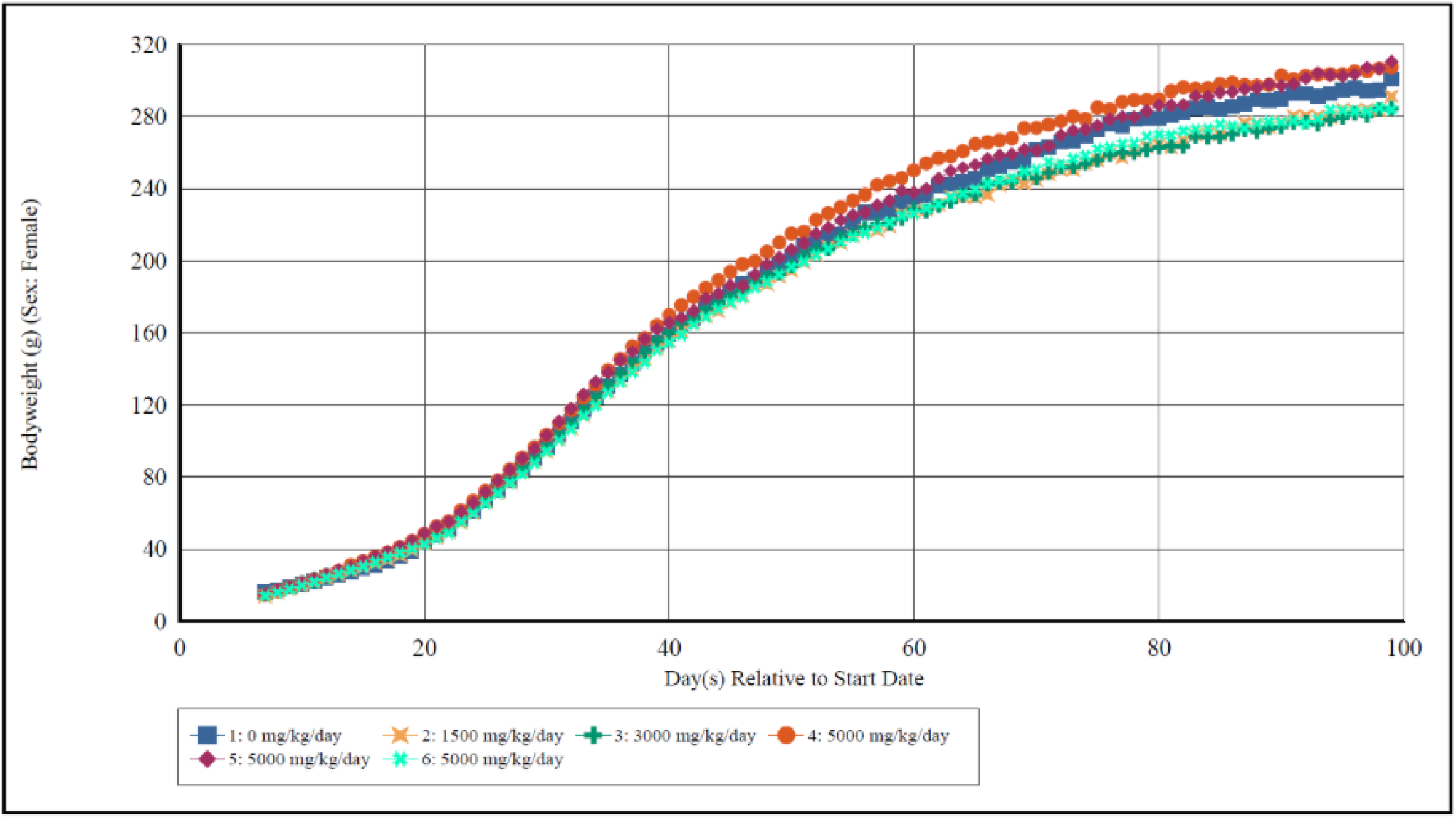
B. Summary of Female Group Mean Body Weight

In the Recovery Phase (PND 127 ± 3), there were no heqLFα-related effects on terminal body weights in either sex. Body weight gains during the Recovery Phase (PND 99–123) were not statistically different from the vehicle control.

Food consumption values at the end of the study were comparable to vehicle controls in both males and females for the Main Study and Recovery Phase. While statistically significant decreases in mean food consumption were observed in females during portions of the Main Study, these differences did not result in changes in terminal body weights and were attributed to the high protein content of the test article rather than to an adverse toxicological effect.

### Developmental and Reproductive Parameters

There were no heqLFα-related effects on preweaning physical development. Although mean days to eye opening showed statistically significant differences at higher dose levels compared with vehicle controls, these findings lacked dose dependency and were similarly observed in the bLF and whey protein comparative control groups, indicating they were not test article-related (**Supplemental Table 1**). All animals evaluated passed auditory startle and pupil constriction assessments on PND 21.

Sexual maturation was not affected by heqLFα treatment. Although some statistically significant differences in the timing and body weight at sexual maturation were observed across dose groups, all values were within the historical control data range, lacked dose dependency, and similar findings were observed in the reference control groups (**Supplemental Table 2**). Thus, these findings were not considered test article-related. Estrous cycle evaluations did not reveal any test article-related changes (**Supplemental Table 3**).

### Neurobehavioral Assessments

Functional observational battery (FOB), motor activity (**Supplemental Table 4**), and Morris water maze assessments (**Supplemental Table 5**) did not demonstrate any heqLFα-related neurobehavioral effects in males or females at any dose level. Performance measures, including latency, sector entries, and trial completion, were comparable to vehicle controls and the bLF and whey protein reference control groups.

### Clinical Pathology

Clinical pathology parameters were evaluated at the end of the Main Study phase (PND 99). Statistically significant alterations were observed in selected hematology (**Table 8**) and clinical chemistry parameters (**Table 9**) in animals administered heqLFα; however, the magnitude of these changes was minimal and lacked associated adverse findings. In hematology assessments, females administered heqLFα at ≥3000 mg/kg bw/day had statistically significant decreases in mean corpuscular hemoglobin (MCH) and corresponding increases in red cell distribution width (RDW) compared with vehicle controls (p ≤ 0.05). In males administered heqLFα at 5000 mg/kg bw/day, similar decreases in MCH and increases in RDW were observed (p ≤ 0.05). Comparable changes were also present in animals administered bLF and whey protein control groups.

**Table 8.** Statistically Significant Effects on Hematological Parameters.

|  |  |  |  |  |  |  |  |  |  |  |
| --- | --- | --- | --- | --- | --- | --- | --- | --- | --- | --- |
| Group | 2 |  | 3 |  | 4 |  | 5 |  | 6 |  |
| Dose (mg/kg bw/day) | 1500 heqLFa |  | 3000 heqLFa |  | 5000 heqLFa |  | 5000 Bovine Lactoferrin |  | 5000 Whey Protein Control |  |
| Sex | M | F | M | F | M | F | M | F | M | F |
| MCH (pg) |  |  |  |  |  |  |  |  |  |  |
| Day 99 | - | - | - | **0.97x** | 0.96x | **0.95x** | **0.96x** | 0.97x | **0.94x** | **0.92x** |
| Day 127 ± 3 | NA | NA | NA | NA | - | - | - | 0.96x | - | - |
| RDW (%) |  |  |  |  |  |  |  |  |  |  |
| Day 99 | - | - | - | **1.08x** | - | - | **1.10x** | 1.05x | **1.07x** | **1.08x** |
| Day 127 ± 3 | NA | NA | NA | NA | - | - | - | - | - | - |

Abbreviations: heqLFa = Human Equivalent Lactoferrin Alpha, M - Males; F - Females.
A dash (—) indicates absence of a test article-related change. **Bolded** values indicate a statistically significant difference from the control group mean. Numerical values are the fold change of the treated group mean value relative to the control group mean value.
NA = Not analyzed.

**Table 9.** Statistically Significant Effects on Clinical Chemistry Parameters.

|  |  |  |  |  |  |  |  |  |  |  |
| --- | --- | --- | --- | --- | --- | --- | --- | --- | --- | --- |
| Group | 2 |  | 3 |  | 4 |  | 5 |  | 6 |  |
| Dose (mg/kg bw/day) | 1500 heqLFα | | 3000 heqLFα | | 5000 heqLFα | | 5000 Bovine Lactoferrin | | 5000 Whey Protein Control | |
| Sex | M | F | M | F | M | F | M | F | M | F |
| Urea Nitrogen (mg/dL) |  |  |  |  |  |  |  |  |  |  |
| Day 99 | - | - | 1.12x | - | 1.18x | - | - | - | - | - |
| Day 127 ± 3 | NA | NA | NA | NA | - | - | - | - | - | - |
| Total Protein (g/dL) |  |  |  |  |  |  |  |  |  |  |
| Day 99 | 0.95x | - | **0.94x** | - | **0.93x** | - | **0.91x** | - | **0.91x** | - |
| Day 127 ± 3 | NA | NA | NA | NA | - | - | - | - | - | - |
| Albumin (g/dL) |  |  |  |  |  |  |  |  |  |  |
| Day 99 | **0.93x** | - | **0.94x** | - | **0.93x** | - | **0.91x** | - | **0.92x** | - |
| Day 127 ± 3 | NA | NA | NA | NA | - | - | - | - | - | - |
| Day 127 ± 3 | NA | NA | NA | NA | - | - | - | - | - | - |
| Calcium (mg/dL) |  |  |  |  |  |  |  |  |  |  |
| Day 99 | 0.97x | - | 0.97x | - | **0.96x** | - | **0.96x** | - | **0.95x** | - |
| Day 127 ± 3 | NA | NA | NA | NA | - | - | - | - | - | - |
| Sodium (mEq/L) |  |  |  |  |  |  |  |  |  |  |
| Day 99 | - | - | **0.97x** | - | **0.97x** | - | - | - | - | - |
| Day 127 ± 3 | NA | NA | NA | NA | - | - | - | - | - | - |
| Chloride (mEq/L) |  |  |  |  |  |  |  |  |  |  |
| Day 99 | - | - | **0.97x** | - | **0.97x** | - | **0.97x** | - | - | - |
| Day 127 ± 3 | NA | NA | NA | NA | - | - | - | - | - | - |

Abbreviations: heqLFα = Human Equivalent Lactoferrin Alpha, M - Males; F - Females.
A dash (—) indicates absence of a test article-related change. **Bolded** values indicate a statistically significant difference from the control group mean. Numerical values are the fold change of the treated group mean value relative to the control group mean value.

In clinical chemistry evaluations, males administered heqLFα at ≥1500 mg/kg bw/day demonstrated decreases in albumin, total protein, and calcium concentrations relative to vehicle controls (p ≤ 0.05). Additionally, males administered ≥3000 mg/kg bw/day showed decreases in sodium and chloride concentrations (p ≤ 0.05), along with increases in urea nitrogen concentrations (p ≤ 0.05). These changes were small in magnitude, remained within or near historical control ranges, and were not associated with correlating histopathological or functional findings.

No heqLFα-related effects were observed on coagulation parameters in either sex at any dose level.

All statistically significant clinical pathology changes observed during the Main Study phase were fully reversed following the Recovery Phase in animals administered heqLFα at 5000 mg/kg bw/day.

### Immunophenotyping

Spleen weight, total splenocyte recovery, and calculated spleen cellularity were not significantly different in animals administered heqLFα compared with vehicle controls in either sex (**Table 10**). A modest but significant decrease in spleen cell viability was observed in female rats administered 3000 mg/kg bw/day heqLFα (*p* < 0.05) and in male rats administered 3000 and 5000 mg/kg bw/day heqLFα (*p* < 0.05) compared with vehicle controls.

**Table 10.**
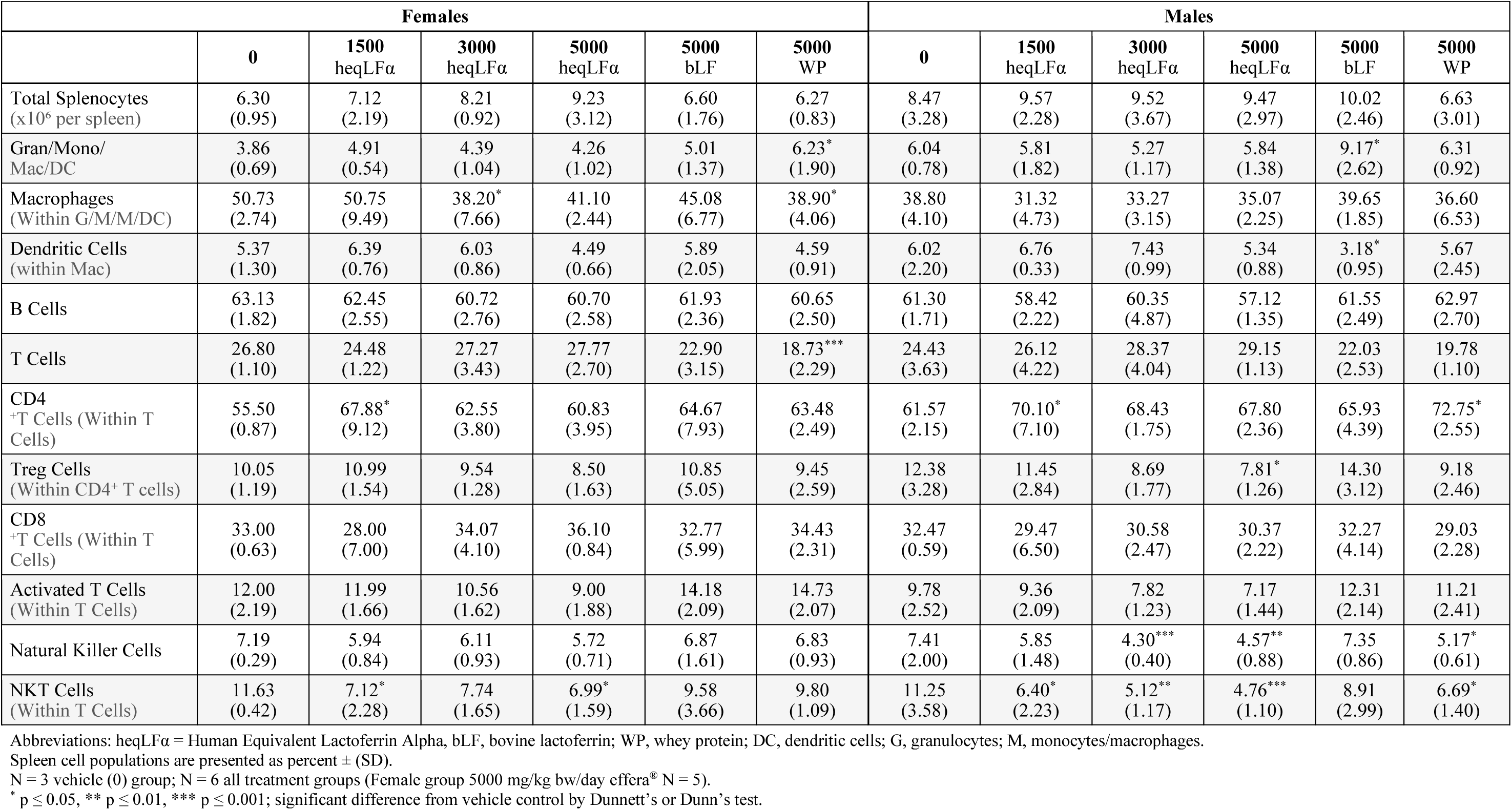
Immunophenotyping Results in Male and Female Rat Spleen Following Oral Gavage Administration of effera^®^, whey protein, or bovine lactoferrin.

Administration of heqLFα at any dose level did not produce statistically significant changes in the percentage of CD172a^+^ myeloid-derived immune cells compared with vehicle control in either sex or in CD172a^+^ CD11b^+^ CD25^+^ dendritic cells. However, a significant reduction in CD172a^+^ CD11b^+^ macrophages was observed in female rats administered 3000 mg/kg bw/day heqLFα (*p* < 0.05) compared with vehicle controls. A similar statistically significant reduction was observed in females administered whey protein (5000 mg/kg bw/day; *p* < 0.05) (**Table 10**).

The frequency of total CD3^+^ T cells was also not significantly altered by heqLFα in either sex. A significant increase in CD4^+^ helper T cells was observed in both sexes administered 1500 mg/kg bw/day heqLFα (*p* < 0.05) relative to vehicle controls. Although non-significant increases were observed at higher dose levels, these did not reach statistical significance. Male rats administered whey protein (5000 mg/kg bw/day) also demonstrated a statistically significant increase in CD4^+^ helper T cells (*p* < 0.01). A moderate decrease in regulatory T cells (CD4^+^ CD25^+^ FoxP3^+^) was observed only in male rats administered 5000 mg/kg bw/day heqLFα (*p* < 0.05) compared with vehicle controls (**Table 10**).

Natural killer (NK) cell percentages were not significantly altered in female rats administered heqLFα. In male rats, statistically significant reductions in NK cells were observed at 3000 mg/kg bw/day (*p* < 0.001) and 5000 mg/kg bw/day (*p* < 0.01) heqLFα compared with vehicle controls. A similar significant reduction was observed in male rats administered whey protein (5000 mg/kg bw/day; *p* < 0.05). Natural killer T (NKT) cell percentages were significantly reduced in both sexes administered heqLFα compared with vehicle controls. In females, reductions were observed at 1500 mg/kg bw/day (*p* < 0.05) and 5000 mg/kg bw/day (*p* < 0.05). In males, statistically significant reductions were observed at 1500 mg/kg bw/day (*p* < 0.05), 3000 mg/kg bw/day (*p* < 0.01), and 5000 mg/kg bw/day (*p* < 0.001). A significant reduction in NKT cells was also observed in male rats administered whey protein (5000 mg/kg bw/day; *p* < 0.05) (**Table 10**).

### Organ Weights

In the kidneys, heqLFα-related statistically significant, minor increases in absolute and/or relative (to brain and/or body weight), compared to the vehicle control, were observed at ≥ 1500 mg/kg bw/day in the males and 5000 mg/kg bw/day in the females (**Table 11**). Compared to the Whey protein control and the historical control data (HCD), absolute and relative to brain kidney weights were increased at the same dose levels in the males and females; however, relative kidney weights to body weight did not differ.

**Table 11.** Summary of Organ Weight (Kidney) Data (Main Study PND 99)

| Table 11. Summary of Organ Weight (Kidney) Data (Main Study PND 99) |  |  |  |  |  |  |  |
| --- | --- | --- | --- | --- | --- | --- | --- |
| | Group 1<br>0 mg/kg/day<br>Vehicle<br>Control<br>(n=10) | Group 2<br>1500<br>mg/kg/day<br>heqLFα<br>(n=10) | Group 3<br>3000 mg/kg/day<br>heqLFα<br>(n=10) | Group 4<br>5000 mg/kg/day<br>heqLFα<br>(n=10) | Group 5<br>5000<br>mg/kg/day<br>Bovine<br>Lactoferrin<br>(Reference<br>Control) | Group 6<br>5000 mg/kg/day<br>Whey Protein<br>Control<br>(n=10) | Historical<br>Control (HCD)<sup>a</sup><br>Mean (g) ±SD<br>(range) |
| **Males** |  |  |  |  |  |  |  |
| No. Animals per Group | 10 | 10 | 10 | 10 | 10 | 10 | - |
| Kidney (No. Weighed) | 10 | 10 | 10 | 10 | 10 | 10 | - |
| Absolute value<br>(% difference from vehicle control) | - | 14.1 | **20.9** | **23.8** | 6.1 | 3.3 | - |
| Absolute value (g) ± SD | 3.3 ± 0.4 | 3.8 ± 0.6 | **4.0\*\* ± 0.5** | **4.1\*\* ± 0.3** | 3.5 ± 0.4 | 3.4 ± 0.4 | 3.1 ± 0.5<br>(2.239–4.140) |
| % of brain weight<br>(% difference from vehicle control) | - | 12.8 | **19.5** | **21.2** | 7.6 | 5.7 | - |
| Absolute percent to brain weight ± SD | 155.5 ± 19.6 | 175.4 ± 26.1 | **185.8\*\* ± 23.4** | **188.4\*\* ± 10.5** | 167.2 ± 16.5 | 164.3 ± 18.6 | 143.4 ± 23.4<br>(105.6–190.3) |
| % of body weight<br>(% difference from vehicle control) | - | **12.2** | **14.8** | **23.6** | 8.3 | 9.9 | - |
| Absolute percent to body weight ± SD | 0.6 ± 0.05 | **0.7\* ± 0.1** | **0.7\* ± 0.08** | **0.8\*\* ± 0.04** | 0.7 ± 0.06 | 0.7 ± 0.03 | 0.6 ± 0.1<br>(0.509–0.844) |
| **Females** |  |  |  |  |  |  |  |
| No. Animals per Group | 10 | 10 | 10 | 10 | 10 | 10 | - |
| Kidney (No. Weighed) | 10 | 10 | 10 | 10 | 9 | 10 | - |
| Absolute value<br>(% difference from vehicle control) | - | 0.8 | 3.0 | **25.1** | **13.3** | 0.8 | - |
| Absolute value (g) ± SD | 1.9 ± 0.2 | 1.9 ± 0.1 | 1.9 ± 0.2 | **2.3\*\* ± 0.2** | **2.1\* ± 0.2** | 1.9 ± 0.3 | 1.8 ± 0.2<br>(1.473–2.334) |
| % of brain weight<br>(% difference from vehicle control) | - | -2.3 | 1.5 | **19.0** | 8.9 | 1.8 | - |
| Absolute percent to brain weight ± SD | 96.2 ± 13.1 | 94.0 ± 5.2 | 97.7 ± 10.1 | **114.5\*\* ± 8.5** | 104.8 ± 9.0 | 97.9 ± 13.1 | 94.1 ± 11.5<br>(74.3–118.2) |
| % of body weight<br>(% difference from vehicle control) | - | 5.3 | 10.1 | **21.2** | 10.8 | 9.4 | - |
| Absolute percent to body weight ± SD | 0.6 ± 0.04 | 0.6 ± 0.03 | 0.7 ± 0.06 | **0.7\*\* ± 0.06** | 0.7 ± 0.06 | 0.7 ± 0.09 | 0.7 ± 0.1<br>(0.559–0.867) |

Abbreviations: heqLFα = Human Equivalent Lactoferrin Alpha, BID = Twice Daily; HCD = historical control data; SD = standard deviation; - = not applicable. Based upon statistical analysis of group means, values highlighted in **bold** are significantly different from vehicle control group – \*P ≤ 0.05, \*\*P ≤ 0.01. <sup>a</sup> HCD compiled from 42 studies conducted in Sprague-Dawley rats (N = 493 (absolute); 492 (relative to body, relative to brain) males and N = 471 (absolute, relative to brain); 470 (relative to body) females), 14-22 weeks of age at necropsy, from 2020 – 2024.

Statistically significant, increased absolute kidney weights, compared to vehicle control, were also observed in the 5000 mg/kg bw/day Bovine Lactoferrin (reference control) in the females.

The decreased absolute and/or relative spleen weights (to brain and/or body weights), compared to vehicle control, were statistically significant in the females at ≥ 1500 mg/kg bw/day heqLFα and 5000 mg/kg bw/day Whey Protein Control group, however, there were no macroscopic or microscopic correlates. These changes lacked dose dependency, and individual weights (compared to the vehicle control) did not demonstrate any pattern or trend; therefore, the weights were not considered related to the administration of heqLFα or Whey Protein Control.

No other test article-related organ weight changes were noted. There were other isolated organ weight values that were statistically different from the respective vehicle control. There were, however, no patterns, trends, or correlating data to suggest these values were relevant; therefore, other organ weight differences observed were considered incidental and unrelated to administration of heqLFα, Bovine Lactoferrin (reference control), or Whey Protein Control.

In the Recovery Phase at PND 127, minor statistically significant increases in absolute kidney weights persisted in males administered 5000 mg/kg bw/day (**Table 12**). Compared to the Whey protein control and the HCD, absolute and relative to brain kidney weights were increased at the same dose level in the males; however, relative kidney weights to body weight did not differ. Statistically significant, increased absolute and/or relative (to brain weight) kidney weights, compared to vehicle control, were also observed in the 5000 mg/kg bw/day Bovine Lactoferrin (reference control) and Whey Protein Control in the males and 5000 mg/kg bw/day Bovine Lactoferrin (reference control) in the females.

**Table 12.** Summary of Organ Weight (Kidney) Data (Recovery Phase PND 127±3)

| Table 12. Summary of Organ Weight (Kidney) Data (Recovery Phase PND 127±3) |  |  |  |  |  |  |  |  |  |  |
| --- | --- | --- | --- | --- | --- | --- | --- | --- | --- | --- |
|  | Males |  |  |  |  | Females |  |  |  |  |
| Group | 1 | 4 | 5 | 6 | HCD<sup>a</sup><br>Mean (g)<br>±SD | 1 | 4 | 5 | 6 | HCD<sup>a</sup><br>Mean (g)<br>±SD |
| Total Dose Level (mg/kg bw/day) | 0 | 5000 | 5000 | 5000 | - | 0 | 5000 | 5000 | 5000 | - |
| BID Dose Level (mg/kg/dose) | 0 | 2500 | 2500 | 2500 | - | 0 | 2500 | 2500 | 2500 | - |
| Test Material | Vehicle<br>Control | heqLFα | Bovine<br>Lactoferrin<br>(ref control) | Whey<br>Protein<br>Control | - | Vehicle<br>Control | heqLFα | Bovine<br>Lactoferrin<br>(ref control) | Whey<br>Protein<br>Control | - |
| No. Animals per Group | 4 | 5 | 5 | 5 | - | 5 | 5 | 3 | 5 | - |
| Kidney (No. Weighed) | 4 | 5 | 5 | 5 | - | 5 | 5 | 3 | 5 | - |
| Absolute value<br>(% difference from vehicle control) | - | **27.4** | **25.6** | **22.9** | - | - | 6.5 | 13.6 | 6.2 | - |
| Absolute value (g) ± SD | 3.1 ± 0.1 | **4.0\*\* ± 0.2** | **3.9\*\* ± 0.2** | **3.8\*\* ± 0.3** | 3.1 ± 0.5<br>(2.239–4.140) | 1.9 ± 0.1 | 2.1 ± 0.1 | 2.2 ± 0.2 | 2.0 ± 0.2 | 1.8 ± 0.2<br>(1.473–2.334) |
| % of brain weight<br>(% difference from vehicle control) | - | **23.9** | **22.1** | 13.8 | - | - | 5.5 | **16.0** | 1.8 | - |
| Absolute percent to brain weight ± SD | 149.8 ± 11.1 | **185.7\*\* ± 15.7** | **182.9\*\* ± 8.2** | 170.5 ± 14.1 | 143.4 ± 23.4<br>(105.6–190.3) | 98.6 ± 7.3 | 104.0 ± 4.6 | **114.4\* ± 9.9** | 100.3 ± 7.3 | 94.1 ± 11.5<br>(74.3–118.2) |
| % of body weight<br>(% difference from vehicle control) | - | **22.5** | 6.5 | 9.6 | - | - | 6.9 | 8.9 | 6.5 | - |
| Absolute percent to body weight ± SD | 0.6 ± 0.03 | **0.7\*\* ± 0.06** | 0.6 ± 0.04 | 0.6 ± 0.02 | 0.6 ± 0.1<br>(0.509–0.844) | 0.6 ± 0.03 | 0.7 ± 0.03 | 0.7 ± 0.05 | 0.7 ± 0.06 | 0.7 ± 0.1<br>(0.559–0.867) |

Abbreviations: heqLFα = Human Equivalent Lactoferrin Alpha, BID = Twice Daily; HCD = historical control data; SD = standard deviation; - = not applicable. Based upon statistical analysis of group means, values highlighted in **bold** are significantly different from vehicle control group – \*P ≤ 0.05, \*\*P ≤ 0.01. <sup>a</sup> HCD compiled from 42 studies conducted in Sprague-Dawley rats (N = 493 (absolute); 492 (relative to body, relative to brain) males and N = 471 (absolute, relative to brain); 470 (relative to body) females), 14-22 weeks of age at necropsy, from 2020 – 2024.

### Macroscopic/Microscopic Observations

There were no heqLFα-related macroscopic findings at scheduled necropsy in either the Main Study or Recovery Phases.

In the kidneys, heqLFα-related increased incidence, and/or severity, compared to the concurrent vehicle control, of tubular mineralization (primarily of tubules at the cortico-medullary junction), tubular basophilia, and/or chronic progressive nephropathy were observed in males and females at ≥ 1500 mg/kg bw/day (**Table 13**). Compared to the Whey protein control, the incidence of the tubular mineralization and basophilia in the heqLFα males was comparable with an increased incidence of the chronic progressive nephropathy and in the heqLFα females the incidence and severity of all three findings were increased. These findings correlated with increased kidney weights at ≥ 1500 mg/kg bw/day in the males and 5000 mg/kg bw/day in the females. The changes in the kidneys were not considered adverse as the findings were minimal to mild, there were no changes in kidney biomarkers, and these findings are routinely observed in Sprague-Dawley rats microscopically.^50^ HCD reveal tubular mineralization in 0.6% males and 7.5% females, tubular basophilia in 16.3% males and 4% females, and chronic progressive nephropathy in 11.7% males and 3.5% females.

**Table 13.**
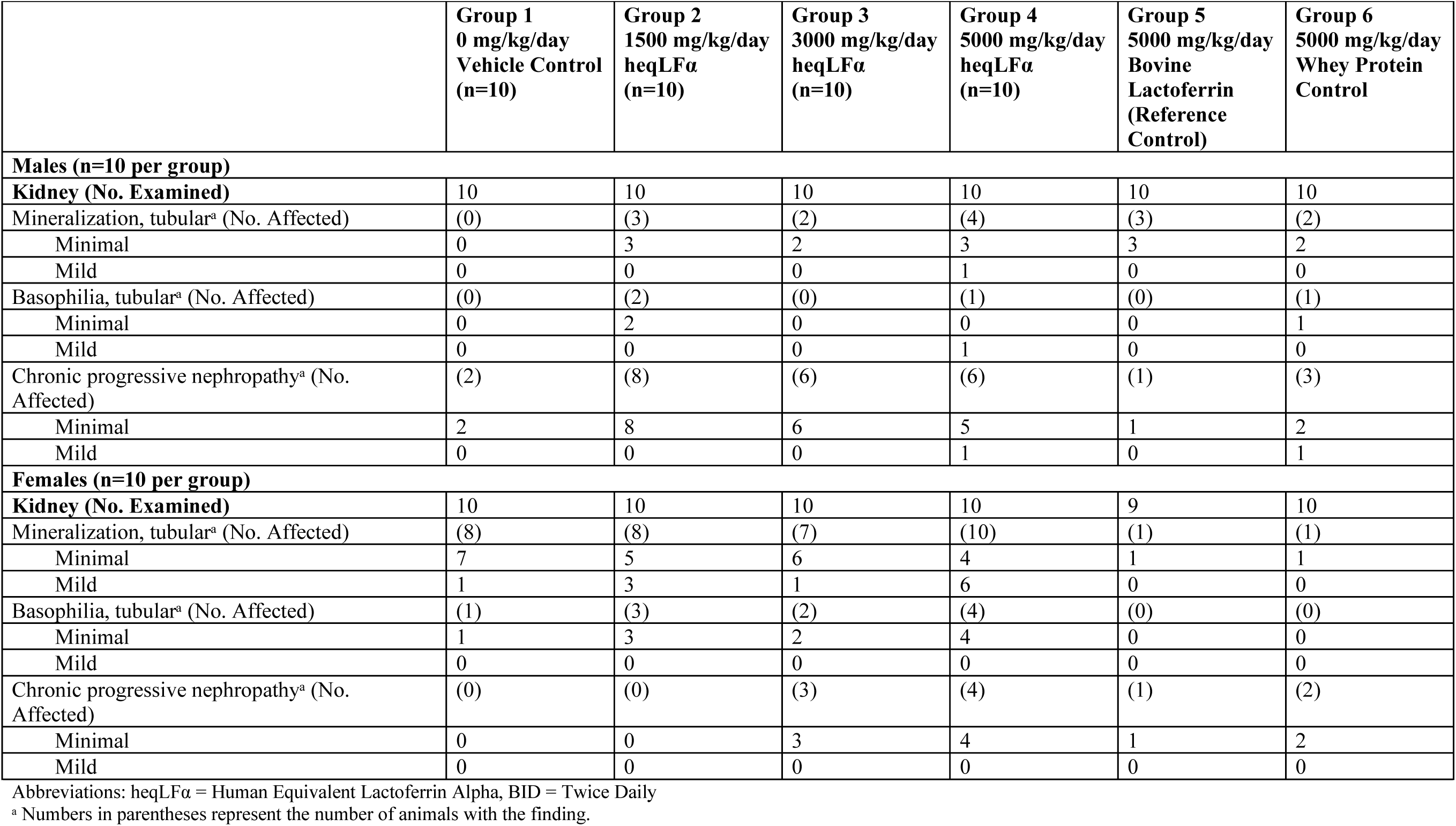
Summary of Microscopic Observations (Main Study, PND 99±3)

In the Recovery Phase, heqLFα-related increased incidence, and/or severity, compared to the concurrent vehicle control, of tubular mineralization was observed in males and females at 5000 mg/kg bw/day and correlated with increased kidney weights in the males (**Table 14**). As heqLFα-related findings of tubular basophilia and chronic progressive nephropathy were not observed, the test article-related findings were considered almost completely recovered and not adverse.

**Table 14.** Summary of Microscopic Observations – Recovery Phase (PND 127±3)

| **Table 14. Summary of Microscopic Observations – Recovery Phase (PND 127±3)** |  |  |  |  |  |  |  |  |
| --- | --- | --- | --- | --- | --- | --- | --- | --- |
|  | **Males** |  |  |  | **Females** |  |  |  |
|  | **Group 1**<br>**0 mg/kg/day**<br>**Vehicle**<br>**Control**<br>**(n=4)** | **Group 4**<br>**5000**<br>**mg/kg/day**<br>**heqLFα**<br>**(n=5)** | **Group 5**<br>**5000**<br>**mg/kg/day**<br>**Bovine**<br>**Lactoferrin**<br>**(n=5)** | **Group 6**<br>**5000**<br>**mg/kg/day**<br>**Whey**<br>**Protein**<br>**Control**<br>**(n=5)** | **Group 1**<br>**0 mg/kg/day**<br>**Vehicle**<br>**Control**<br>**(n=5)** | **Group 4**<br>**5000**<br>**mg/kg/day**<br>**heqLFα**<br>**(n=5)** | **Group 5**<br>**5000**<br>**mg/kg/day**<br>**Bovine**<br>**Lactoferrin**<br>**(n=3)** | **Group 6**<br>**5000**<br>**mg/kg/day**<br>**Whey**<br>**Protein**<br>**Control**<br>**(n=5)** |
| **Kidney (No. Examined)** | 4 | 5 | 5 | 5 | 5 | 5 | 3 | 5 |
| Mineralization, tubular<sup>a</sup> (No. Affected) | (0) | (3) | (2) | (2) | (1) | (4) | (0) | (4) |
| Minimal | 0 | 3 | 2 | 2 | 1 | 2 | 0 | 4 |
| Mild | 0 | 0 | 0 | 0 | 0 | 2 | 0 | 0 |

Abbreviations: heqLFα = Human Equivalent Lactoferrin Alpha, BID = Twice Daily; Groups 2 and 3 were not included in the Recovery Phase.
<sup>a</sup> Numbers in parentheses represent the number of animals with the finding.

Other microscopic findings observed were considered incidental, of the nature commonly observed in this strain and age of rats, and/or were of similar incidence and severity in the vehicle control and treated animals and, therefore, were considered unrelated to administration of heqLFα.

### Toxicokinetics

Serum heqLFα concentrations were generally below the limit of quantification in control animals, with minor exceptions in two animals on PND 98, which did not impact the study interpretation. Systemic exposure to heqLFα following twice-daily oral gavage administration was confirmed in juvenile male and female rats at dose levels of 1500, 3000, and 5000 mg/kg bw/day with no apparent sex differences in exposure. Serum concentrations of heqLFα were detectable at the earliest post-dose sampling time point, indicating rapid absorption following oral administration (**Table 15**).

**Table 15.** Toxicokinetic Parameters of heqLFα in Male and Female Juvenile Rat Serum Following Oral Gavage Administration of heqLFα.

| PND | Sex | Dose (mg/kg/day) | C<sub>max</sub> (ng/mL) | t<sub>max</sub> (hr) | t<sub>last</sub> (hr) | AUC<sub>0-24h</sub> | F:M AUC<sub>0-24h</sub> | F:M C<sub>max</sub> | R<sub>AUC</sub> | RC<sub>max</sub> |
| --- | --- | --- | --- | --- | --- | --- | --- | --- | --- | --- |
| 7 | Combined | 1500 | 243000 | 8.5 | 24 | 669000 | NA<sup>a</sup> | NA | NA | NA |
|  |  | 3000 | 589000 | 8.5 | 24 | 1620000 | NA<sup>a</sup> | NA | NA | NA |
|  |  | 5000 | 329000 | 8.5 | 24 | 911000 | NA<sup>a</sup> | NA | NA | NA |
|  | Female | 1500 | 2030 | 8.5 | 24 | 5860 | 0.00440 | 0.00419 | NA | NA |
|  |  | 3000 | 252 | 8.5 | 24 | 1890 | 0.000584 | 0.000214 | NA | NA |
|  |  | 5000 | 7790 | 8.5 | 24 | 24000 | 0.0133 | 0.0120 | NA | NA |
|  | Male | 1500 | 484000 | 8.5 | 24 | 1330000 | NA<sup>a</sup> | NA | NA | NA |
|  |  | 3000 | 1180000 | 8.5 | 24 | 3240000 | NA<sup>a</sup> | NA | NA | NA |
|  |  | 5000 | 651000 | 8.5 | 24 | 1800000 | NA<sup>a</sup> | NA | NA | NA |
| 98 | Combined | 1500 | 93.6 | 0.5 | 24 | 260 | NA<sup>a</sup> | NA | 0.000388 | 0.000385 |
|  |  | 3000 | 41.3 | 0.5 | 12 | NA<sup>a</sup> | NA<sup>a</sup> | NA | 5.83E-05 | 7.02E-05 |
|  |  | 5000 | 7080 | 0.5 | 24 | 6750 | NA<sup>a</sup> | NA | 0.00741 | 0.0215 |
|  | Female | 1500 | 152 | 0.5 | 12 | NA<sup>a</sup> | 2.73 | 17.5 | 0.0453 | 0.0748 |
|  |  | 3000 | 73.8 | 0.5 | 12 | 117 | 0.980 | 5.92 | 0.0485 | 0.293 |
|  |  | 5000 | 234 | 12 | 24 | 1880 | 0.162 | 0.0166 | 0.0785 | 0.0301 |
|  | Male | 1500 | 8.69 | 1.5 | 24 | 97.3 | NA<sup>a</sup> | NA | 7.30E-05 | 1.79E-05 |
|  |  | 3000 | 12.5 | 12 | 12 | NA<sup>a</sup> | NA<sup>a</sup> | NA | 2.89E-05 | 1.06E-05 |
|  |  | 5000 | 14100 | 0.5 | 24 | 11600 | NA<sup>a</sup> | NA | 0.00646 | 0.0217 |

Abbreviations: heqLFα = human equivalent lactoferrin alpha; PND = postnatal day, F:M = female-to-male ratio, NA = not applicable; R<sub>AUC</sub>, AUC<sub>t<sub>last</sub></sub> PND 98 / AUC<sub>t<sub>last</sub></sub> PND 7, RC<sub>max</sub>, C<sub>max</sub> PND 98 / C<sub>max</sub> PND 7.
<sup>a</sup>: AUC<sub>0-24h</sub> was not reportable because %AUC extrapolation exceeded 25% of total area

Mean concentration–time profiles demonstrated dose-related increases in maximum observed concentration (C_max_) and area under the concentration–time curve to the last quantifiable time point (AUC_tlast_) on both the initial toxicokinetic assessment day (PND 7) and following repeated administration (PND 98). Detectable serum concentrations generally declined over the sampling interval, consistent with systemic clearance following oral exposure (**Table 15**).

At the no observed adverse effect level (NOAEL) dose of 5000 mg/kg bw/day, mean C_max_ and AUC_tlast_ values (combined sexes) following 13 weeks of dosing were approximately 7080 ng/mL and 6750 hr*ng/mL, respectively. Systemic exposure appeared lower at the later sampling interval and showed no accumulation. Evaluation of accumulation following repeated administration did not demonstrate disproportionate systemic accumulation across dose groups. Exposure parameters on PND 98 were generally comparable to or lower than those observed at earlier time points, indicating no evidence of progressive systemic accumulation over the dosing period (**Table 15**).

## Discussion/Conclusion

LF is broadly associated with improved infant health and is therefore considered an appealing food ingredient to be added to infant formula and products for babies and children but must first be thoroughly safety tested. Here, we show that heqLFα has been thoroughly evaluated to indicate that 1) heqLFα has no genotoxic risk potential and 2) heqLFα is safe and well-tolerated in juvenile rats at doses up to 5000 mg/kg bw/day following twice-daily administration for 90 days, starting at PND7, the equivalent of birth in humans. No functional renal impairment, developmental toxicity, neurobehavioral changes, or clinically meaningful immunotoxic effects were identified. The NOAEL of 5000 mg/kg bw/day, expressed on a pure heqLFα basis, corresponds to approximately a 9.2-fold margin of exposure relative to the estimated high-end infant formula intake of 546 mg/kg bw/day (based on EFSA reference consumption of 260 mL/kg bw/day and a 2.1 g/L use level).^34^ Collectively, these findings support the safety of heqLFα under intended conditions of use, including use in infant populations.

The absence of mutagenic or clastogenic activity observed with heqLFα aligns with published genotoxicity data for HiMOs^36,37^ and bLF.^51^ Similarly, another milk bioactive protein, bovine osteopontin (bOPN), has been shown to lack mutagenic or genotoxic activity in Ames assays and chromosomal damage models.^52^ These data support the conclusion that major human milk bioactives including oligosaccharides and glycoproteins show no evidence of genotoxicity potential.

To date, the majority of repeated-dose toxicology studies evaluating milk bioactives have been conducted in young adult rats (typically 4–7 weeks of age at study initiation). Representative 28-and 90-day studies of hLF from bioengineered sources or milk-derived LF preparations were initiated in 4–6-week-old rats,^21–24,35^ and the recent heqLFα 4-week immunotoxicity study similarly used 6-week-old rats^19^. These studies consistently demonstrated NOAELs at the top doses tested with no toxicologically relevant findings but they do not specifically model the physiological immaturity and rapid organ development characteristic of early infancy. ^53^

Both bLF^12,13^ and bOPN^54^ have been approved for use in infant formula in the EU following comprehensive safety evaluations, and bLF is also permitted for use in the United States; however, these assessments relied largely on subchronic studies conducted in young adult rodents. The current EFSA guidance for substances intended for use in foods for infants younger than 16 weeks of age recognizes that standard adult rodent studies are insufficient to fully characterize risk for this vulnerable population and recommends study designs that incorporate neonatal exposure and developmental endpoints.^34^ An Institute of Medicine (IOM) report also emphasizes using appropriate animal models at relevant developmental stages.^55^

The present 13-week GLP juvenile rat study with heqLFα meets the current EFSA guidance and IOM recommendations, as it was initiated during early postnatal life and spans critical windows of organ development including renal, immune, and neurological systems. This study represents the most comprehensive nonclinical safety evaluation to date of heqLFα, incorporating repeated-dose toxicity from PND 7 through PND 98, toxicokinetics, immunophenotyping, and a Recovery Phase. There were no toxicologically meaningful heqLFα-related adverse effects following twice-daily oral gavage at doses up to 5000 mg/kg bw/day compared to saline vehicle control.

Mean body weights and body weight gains in both male and female rats administered heqLFα at 1500, 3000, or 5000 mg/kg bw/day remained within normal ranges for age and sex and were comparable to vehicle and protein control groups, indicating no adverse impact on growth or development.

No heqLFα-related adverse clinical pathology findings were observed. Statistically significant changes compared to the vehicle control were either comparable in bLF and/or whey protein, consistent with a high protein diet, or were small in magnitude, remained within or near historical control ranges, and were not associated with correlating histopathological or functional findings.

Modest treatment-related changes were observed in spleen-resident immune cell populations in male and female rats; however, these effects were often observed concurrently in the whey protein control group, suggesting that the changes are not due to biological activity specific to heqLFα itself. Therefore, based solely on the immunophenotyping analysis conducted in this study, there is no evidence of immunotoxicity that can be specifically ascribed to the biological activity of heqLFα.

Increased kidney weights were observed in males at ≥1500 mg/kg bw/day and in females at 5000 mg/kg bw/day. Importantly, these changes were not accompanied by adverse alterations in clinical pathology biomarkers that would indicate renal dysfunction. The absence of functional impairment suggests that the kidney weight increases represent adaptive hypertrophy rather than adverse toxicity. This interpretation is consistent with the well-documented renal response to high dietary protein intake in rodents, wherein increased renal workload results in compensatory enlargement without structural degeneration.^53,56–58^ The similarity of findings in the whey protein control group further supports a protein-mediated physiological adaptation rather than a lactoferrin-specific toxicologic effect.

The microscopic renal findings observed in this study (tubular mineralization, basophilia, and chronic progressive nephropathy (CPN)) are also well-characterized background lesions in Sprague-Dawley rats.^50,59^ Tubular mineralization and tubular basophilia are commonly reported spontaneous or diet-associated findings, particularly in rapidly growing rodents or those exposed to increased protein or electrolyte load.^50^ CPN is a strain-related, age-dependent lesion in rats and is widely recognized as having limited relevance to humans.^59^ The minimal-to-mild severity, absence of progressive worsening during Recovery Phase, and lack of correlating functional impairment support the conclusion that these lesions are non-adverse and consistent with known background pathology in this strain. Systemic exposure (C_max_ and AUC) decreased between PND 7 and PND 98. This reduction in systemic exposure over time further argues against cumulative nephrotoxicity. The Recovery Phase demonstrated resolution of tubular basophilia and CPN findings and near-complete resolution of mineralization, reinforcing the reversibility and adaptive nature of the renal observations.

Systemic exposure to heqLFα tended to increase with dose, although high inter-animal variability was observed. Systemic exposure was lower on PND 98 compared to PND 7, with C_max_ values ranging from 7.02E-05 to 0.0215 and accumulation values less than 1 (5.83E-05 to 0.00741), indicating no accumulation. Mean heqLFα was quantifiable in rats up to 24 hours post first daily dose at all dose levels on PND 7 and up to 12 or 24 hours post first daily dose on PND 98. t_max_ values for heqLFα were observed by 8.5 hours post first daily dose (0.5-hour post 2nd daily dose) on PND 7 and by 0.5-hour post first daily dose on PND 98. These TK findings are consistent with those observed in adult rats in the 28-day GLP study of the same ingredient,^19^ in which heqLFα demonstrated low systemic bioavailability following oral administration, rapid detection in serum, substantial inter-animal variability, and no evidence of accumulation with repeated dosing. In both adult and juvenile models, systemic exposure decreased over time and was not associated with adverse clinical, pathological, or immunological findings. Collectively, these data show no exposure–response relationship with toxicity endpoints, which supports the overall safety profile of heqLFα across the lifecycle.

In conclusion, no heqLFα-related adverse effects were observed for survival, body weight, food consumption, clinical observations, development or neurobehavioral assessments, hematology, coagulation, clinical chemistry, toxicokinetics, immunophenotyping, or histopathology in male and female Sprague-Dawley rats administered heqLFα twice daily for 90 days beginning at PND 7 compared to a saline vehicle control. This study further demonstrates that heqLFα intake of up to 5000 mg/kg bw/day for 90 days in a juvenile rat model is comparable overall in safety and toxicological endpoints to bLF and whey protein and is safe and well tolerated with a NOAEL of 5000 mg/kg bw/day, the highest dose evaluated.

