## Supplemental Tables for "Preclinical Safety Evaluation of Human Lactoferrin Alpha (Effera^®^)"

**Supplemental Table 1. Summary of Reflex and Development**

| **Endpoint** | **Statistic** | **0 mg/kg bw /day Vehicle Control** | **1500 mg/kg bw /day heqLFα** | **3000 mg/kg bw/day heqLFα** | **5000 mg/kg bw/day heqLFα** | **5000 mg/kg bw/day Bovine LF** | **5000 mg/kg bw/day Whey Protein** |
|---|---|---|---|---|---|---|---|
| **Males** | | | | | | | |
| Day [k] Eyes Opened | Mean | 13.4 | 14.7 | $16.0^{**}$ | $15.3^{**}$ | $15.8^{**}$ | $15.6^{**}$ |
| | SD | 1.1 | 0.7 | 0.0 | 1.0 | 0.5 | 0.5 |
| | N | 12 | 9 | 4 | 8 | 4 | 10 |
| | %Diff | – | 9.3 | 19.3 | 13.7 | 17.4 | 16.3 |
| Day Pupil [k] Constriction | Mean | 21.0 | 21.0 | 21.0 | 21.0 | 21.0 | 21.0 |
| | SD | 0.0 | 0.0 | 0.0 | 0.0 | 0.0 | 0.0 |
| | N | 15 | 10 | 10 | 14 | 13 | 15 |
| | %Diff | – | 0.0 | 0.0 | 0.0 | 0.0 | 0.0 |
| Day Auditory [k] Startle | Mean | 21.0 | 21.0 | 21.0 | 21.0 | 21.0 | 21.0 |
| | SD | 0.0 | 0.0 | 0.0 | 0.0 | 0.0 | 0.0 |
| | N | 15 | 10 | 10 | 14 | 13 | 15 |
| | %Diff | – | 0.0 | 0.0 | 0.0 | 0.0 | 0.0 |
| **Females** | | | | | | | |
| Day [k] Eyes Opened | Mean | 13.5 | 14.8 | $16.0^{**}$ | $15.4^{*}$ | $16.0^{n}$ | $15.5^{**}$ |
| | SD | 1.3 | 0.4 | 0.0 | 0.9 | – | 0.5 |
| | N | 13 | 5 | 4 | 5 | 1 | 10 |
| | %Diff | – | 9.9 | 18.9 | 14.4 | 18.9 | 15.1 |
| Day Pupil [k] Constriction | Mean | 21.0 | 21.0 | 21.0 | 21.0 | 21.0 | 21.0 |
| | SD | 0.0 | 0.0 | 0.0 | 0.0 | 0.0 | 0.0 |
| | N | 15 | 9 | 10 | 12 | 10 | 15 |
| | %Diff | – | 0.0 | 0.0 | 0.0 | 0.0 | 0.0 |
| Day Auditory [k] Startle | Mean | 21.0 | 21.0 | 21.0 | 21.0 | 21.0 | 21.0 |
| | SD | 0.0 | 0.0 | 0.0 | 0.0 | 0.0 | 0.0 |
| | N | 15 | 9 | 10 | 12 | 10 | 15 |
| | %Diff | – | 0.0 | 0.0 | 0.0 | 0.0 | 0.0 |

Abbreviations: heqLFα = Human Equivalent Lactoferrin Alpha. [k] - Kruskal-Wallis & Dunn, 2-Sided: * = $p \leq 0.05$; ** = $p \leq 0.01$

| Supplemental Table 2. Mean Number of Days to Sexual Maturation (Male and Female) | | | | | | | | |
|---|---|---|---|---|---|---|---|---|
| **Sex** | **Sexual Maturation** | **Mean Number of Post Natal Days to Complete** | | | | | | **HCD Range** |
| | | **heqLFα** | | | | **Bovine LF** | **Whey Protein** | |
| | | **0 mg/kg bw/day** | **1500 mg/kg bw/day** | **3000 mg/kg bw/day** | **5000 mg/kg bw/day** | **5000 mg/kg bw/day** | **5000 mg/kg bw/day** | |
| Male | Balano-preputial separation | 45.3 | 45.6 | 46.4 | 44.5 | 42.8 | 45.3 | 40.6 - 47.3 |
| Female | Vaginal Opening | 31.2 | 32.3 | **34.0**[a] | 33.0 | 32.2 | **33.1**[a] | 29.3 – 34.2 |

Abbreviations: heqLFα = Human Equivalent Lactoferrin Alpha, HCD = Historical Control Data

[a] Statistical significance = $p \leq 0.01$

**Supplemental Table 3. Summary of Estrous Cycling: (Day(s) d86→d99)**

| Endpoint | Statistic | 0 mg/kg bw /day Vehicle Control | 1500 mg/kg bw /day heqLFα | 3000 mg/kg bw/day heqLFα | 5000 mg/kg bw/day heqLFα | 5000 mg/kg bw/day Bovine LF | 5000 mg/kg bw/day Whey Protein |
|---|---|---|---|---|---|---|---|
| **Females** | | | | | | | |
| Group Size | | 10 | 10 | 10 | 10 | 9 | 10 |
| Number of Cycles d86→d99 [k] | Mean | 1.9 | 1.9 | 1.8 | 2.3 | 1.7 | 2.3 |
| | SD | 0.7 | 0.3 | 0.4 | 0.5 | 0.9 | 0.7 |
| | N | 10 | 10 | 10 | 10 | 9 | 10 |
| Mean Cycle Lengths (Days) d86→d99 [k] | Mean | 3.95 | 4.00 | 4.55 | 3.82 | 4.44 | 4.15 |
| | SD | 1.00 | 0.24 | 0.69 | 0.74 | 1.47 | 0.51 |
| | N | 10 | 10 | 10 | 10 | 8 | 10 |
| | %Diff | – | 1.27 | 15.19 | -3.38 | 12.34 | 5.06 |

**Summary of Estrous Cycles: (Day(s) d110→d130)**

| Endpoint | Statistic | 0 mg/kg bw /day Vehicle Control | 5000 mg/kg bw/day heqLFα | 5000 mg/kg bw/day Bovine LF | 5000 mg/kg bw/day Whey Protein |
|---|---|---|---|---|---|
| **Females** | | | | | |
| Group Size | | 5 | 5 | 5 | 5 |
| Number of Cycles Recovery Phase d110→d130 [k] | Mean | 2.0 | 2.4 | 2.3 | 2.2 |
| | SD | 0.7 | 0.9 | 0.6 | 0.4 |
| | N | 5 | 5 | 3 | 5 |
| | %Diff | – | 20.0 | 16.7 | 10.0 |
| Mean Cycle Lengths Recovery Phase (Days) d110→d130 [k] | Mean | 4.30 | 3.60 | 3.72 | 4.40 |
| | SD | 0.45 | 0.42 | 0.25 | 0.74 |
| | N | 5 | 5 | 3 | 5 |
| | %Diff | – | -16.28 | -13.44 | 2.33 |

Abbreviations: heqLFα = Human Equivalent Lactoferrin Alpha
[k] – Kruskal-Wallis & Dunn, 2-Sided: No Significance at $p \leq 0.05$

**Supplemental Table 4. Motor Activity Summary**

| Subset | Sex | Session | Motor Activity Data – Combined Trials | | | | | |
|---|---|---|---|---|---|---|---|---|
| | | | heqLFα | | | | Bovine LF | Whey Protein |
| | | | 0 mg/kg bw/day | 1500 mg/kg bw/day | 3000 mg/kg bw/day | 5000 mg/kg bw/day | 5000 mg/kg bw/day | 5000 mg/kg bw/day |
| Main | Male | Ambulation | 124.93 | 109.52 | 119.35 | 89.10 | 126.05 | 179.22 |
| | | Fine Movement | 697.37 | 570.92 | 640.12 | 470.83 | 640.85 | 869.45 |
| | Female | Ambulation | 184.20 | 315.15[a] | 214.35 | 189.58 | 347.24[a] | 229.95 |
| | | Fine Movement | 677.63 | 898.48 | 678.77 | 659.62 | 1032.91[a] | 775.02 |
| Recovery | Male | Ambulation | 158.58 | NA | NA | 94.27 | 85.37 | 62.47 |
| | | Fine Movement | 782.88 | NA | NA | 473.90 | 458.77 | 356.57[a] |
| | Female | Ambulation | 209.97 | NA | NA | 206.23 | 189.56 | 176.83 |
| | | Fine Movement | 791.20 | NA | NA | 678.87 | 625.06 | 604.20 |

Abbreviations: heqLFα = Human Equivalent Lactoferrin Alpha, NA=Not Applicable

[a] Statistical significance $p \leq 0.05$

**Supplemental Table 5. Morris Water Maze Summary**

| Subset | Sex | Session | Mean Latency (sec) to Reach Goal Platform (Combined Trials) | | | | | |
|---|---|---|---|---|---|---|---|---|
| | | | **heqLFα** | | | | **Bovine LF** | **Whey Protein** |
| | | | **0 mg/kg bw/day** | **1500 mg/kg bw/day** | **3000 mg/kg bw/day** | **5000 mg/kg bw/day** | **5000 mg/kg bw/day** | **5000 mg/kg bw/day** |
| Main | Male | 1 | 27.39 | 30.51 | 30.69 | 22.84 | 27.30 | 28.50 |
| | | 2 | 11.99 | 13.36 | 15.23 | 11.57 | 10.64 | 12.79 |
| | Female | 1 | 27.49 | 34.74 | 35.56 | 28.99 | 24.74 | 33.31 |
| | | 2 | 11.80 | 17.08 | 18.99 | 18.14 | 11.42 | 18.38 |
| Recovery | Male | 1 | 22.00 | NA | NA | 28.33 | 25.73 | 39.42 |
| | | 2 | 14.69 | NA | NA | 15.33 | 10.42 | 13.93 |
| | Female | 1 | 28.56 | NA | NA | 28.49 | 36.78 | 34.09 |
| | | 2 | 13.19 | NA | NA | 11.56 | 20.70 | 18.98 |

Abbreviations: heqLFα = Human Equivalent Lactoferrin Alpha, NA=Not Applicable
